# Community-wide adaptive evolution of gut bacteria during dietary transitions

**DOI:** 10.64898/2026.10.01.755998

**Authors:** Samantha G. Hernandez, Siddharth Uppal, Dong-Min Jin, Abigail A. Landers, Nathaniel Ennis, Daniel D. Sprockett, Andrew H. Moeller

## Abstract

Diet exerts selection on bacterial species within the mammalian gut microbiota^1,2,3,4,5^, but the extent, modes, genomic bases, and repeatability of gut bacterial adaptive evolution in response to dietary variation within hosts are poorly understood. Here, we show that within-host dietary transitions drive ubiquitous and highly reproducible adaptive evolution of gut bacterial populations. We deeply profiled allele-frequency dynamics in gut microbiota of an experimental cohort of litter-matched wild-derived isogenic mice either transitioned from low-fat/high-carbohydrate diets to high-fat/low-carbohydrate diets or maintained on their ancestral low-fat/high-carbohydrate diet. Dietary transitions induced significantly parallel allele-frequency changes across replicates in 61/62 core gut bacterial species within five months, whereas 0/62 species displayed significant across-replicate parallelism in the control group, indicating deterministic adaptive evolutionary responses across the gut microbiota. Adaptation proceeded through multiple modes, including strain sorting and gene-wise responses. The latter were enriched in a subset of gene families and functions across bacterial species, including invertases. Culturing revealed an invertible DNA switch in *Phocaeicola sartorii* that, in all replicates transitioned to high-fat/low-carbohydrate diets, localized a promoter sequence upstream of cell-surface glycan foraging genes. These findings elucidate widespread gut bacterial adaptive evolution mediated by within-host dietary variation.

## Main

Mammals harbor complex gut microbial communities whose species can evolve—and potentially adapt—within individual hosts^4,5,6,7,8,9,10,11,12,13,14,15,16^. Changes in selection pressures, such as dietary transitions or antibiotic exposures, have been shown to drive deterministic changes in the relative abundances of gut bacterial species within individual hosts^1,2,17,18^, and selection-induced changes in the microbiota can, in turn, affect host immunity, metabolism, and health^19,20^. Recent retrospective studies have detected signatures of selection in gut bacterial genomes^21,22,23,24^, suggesting histories of within-host adaptive evolution of gut bacterial species. However, resolving the full extent of within-host gut bacterial adaptation has been limited by the difficulty of tracking evolutionary dynamics at strain-, gene-, and allele-level resolution in complex microbiota^25^. Retrospective studies have generally examined genetically heterogeneous hosts exposed to uncontrolled and heterogeneous environmental conditions^21,22,23,24^, in which the absence of a defined perturbation and matched controls has precluded attributing allele frequency change to specific selective regimes. Thus, prior work has been poorly equipped to investigate whether microbiota-wide adaptive responses of gut bacterial populations to specific selective regimes are deterministic and reproducible or stochastic and difficult to predict.

The extent to which evolutionary responses to environmental change are repeatable given similar initial conditions represents a longstanding question in evolutionary biology, but its investigation has been primarily restricted to focal populations, lineages, or traits. Experimental and comparative studies have shown that natural selection can generate similar outcomes across replicate populations and independent lineages^26,27^. At the genetic level, repeated adaptation can likewise involve a restricted and therefore partly predictable set of genes and pathways^28,29^. However, far less is known about the repeatability of evolution at the scale of entire communities, in which environmental change simultaneously imposes selection on many interacting populations. In communities, nonlinear interactions among species can generate dynamical instability and even chaos^30,31,32^, potentially amplifying sensitivity to initial conditions and reducing the predictability of adaptive evolutionary trajectories. Understanding the extent of determinism during adaptive evolution in the context of natural communities is important because species interactions can alter adaptation to novel environments^33,34^ in ways that influence ecosystem functioning^35^.

Evolve-and-resequence (E&R) experiments, in which population-genetic variation is measured before and after exposure to an environment, provide a powerful approach for delineating natural selection from genetic drift and for quantifying the extent of determinism during adaptive evolution^29,36^. However, community-level E&R studies have largely relied on simplified synthetic communities or followed evolution (i.e., changes in the frequencies of genetic variants) in selected focal populations. For example, previous E&R experiments focused on gut microbiota have tracked evolutionary dynamics of individual gut bacterial species (e.g., through experiments with strains genetically engineered with synthetic barcodes or fluorescent markers) or low-complexity (e.g., 12-species) synthetic communities in gnotobiotic laboratory mice^4,5,10,12^. The limited scope of prior experiments has precluded testing whether adaptive evolution is repeatable across the diversity of bacterial populations comprising natural microbiota, given similar initial conditions.

Here, we conducted E&R experiments focused on the microbiota of a wild-derived inbred line of house mice (*Mus musculus domesticus*) originating from a wild breeding pair from Saratoga Springs, NY and inbred in laboratory housing for >45 generations. Unlike standard laboratory mouse lines (e.g., C57BL/6), this wild-derived line has never been rederived under sterile conditions and has retained a relatively diverse microbiota, including endogenous house-mouse gut bacterial lineages that have been lost from laboratory mice since their transition into captivity >120 years ago^37,38,39^. Thus, this wild-derived line enables experimental interrogation of endogenous microbiota constituents within the context of an isogenic host genetic background. To impose a defined selective pressure, we subjected mice to a dietary transition from their standard low-fat/high-carbohydrate diet to a high-fat/low-carbohydrate diet, a perturbation known to alter gut-microbiota composition^40^. We replicated this transition across independent cages while maintaining litter-matched controls on the ancestral diet. Combining deep sequencing and bacterial-isolate cultivation of metagenomic samples collected during these E&R experiments with a *de novo* bespoke reference genome database assembled from experimental mice allowed us to quantify adaptive evolutionary responses across gut bacterial species within a complex mammalian gut microbiota in its native host.

## Experimental evolution of mouse gut microbiota

To interrogate adaptive evolutionary responses of gut bacterial species during dietary transitions, we split mouse littermates into two groups (n=15 mice per diet group; see Methods). The control group was maintained on a low-fat diet (LF) representing the ancestral macronutrient profile on which mice were maintained prior to the experiment. The experimental group was transitioned to a high-fat diet (HF) (Fig. 1a; Extended Data Table 1) (see Methods). Next, we generated genome-resolved metagenomic datasets from samples collected before and during the E&R experiments (7–15 samples per mouse). We sequenced metagenomes from 439 fecal samples collected from individual mice over a 5-month period (Supplementary Table 1), including deep sequencing of samples collected within 1 day before the diet shift (PRE), and 1 week (POST) and 3–5 months (END) after the diet shift (mean = 28,580,244, standard deviation = 6,184,910 reads per sample) (Extended Data Fig. 1). Using these data, we generated 2,218 high-quality bacterial metagenome-assembled genomes (MAGs) (Extended Data Fig. 2) (see Methods), representing 160 species-level clusters (‘species’) based on 95% average nucleotide identity (ANI) (Supplementary Table 2). On average, 79% of high-quality host-filtered reads per metagenome mapped to the 160 dereplicated genomes of these species, indicating that the genome database was representative of mouse metagenomes. Ecological analyses revealed that gut-microbiota species composition (Bray-Curtis dissimilarities) changed in response to HF diets (Fig. 1b; Extended Data Figs. 3–4; Supplementary Results) (PERMANOVA; *p*-values < 0.001 for litter-matched comparisons between HF- and LF-fed mice at both POST and END timepoints; Supplementary Table 3). HF diets reduced microbiota alpha diversity (Shannon, two-tailed Wilcoxon test, Benjamini–Hochberg-corrected *p* = 0.0058) (Supplementary Tables 3 and 4; Extended Data Fig. 5). Together, these results indicate that the HF diet induced deterministic changes in the species composition of gut microbiota across replicated experiments.

**Fig. 1:**
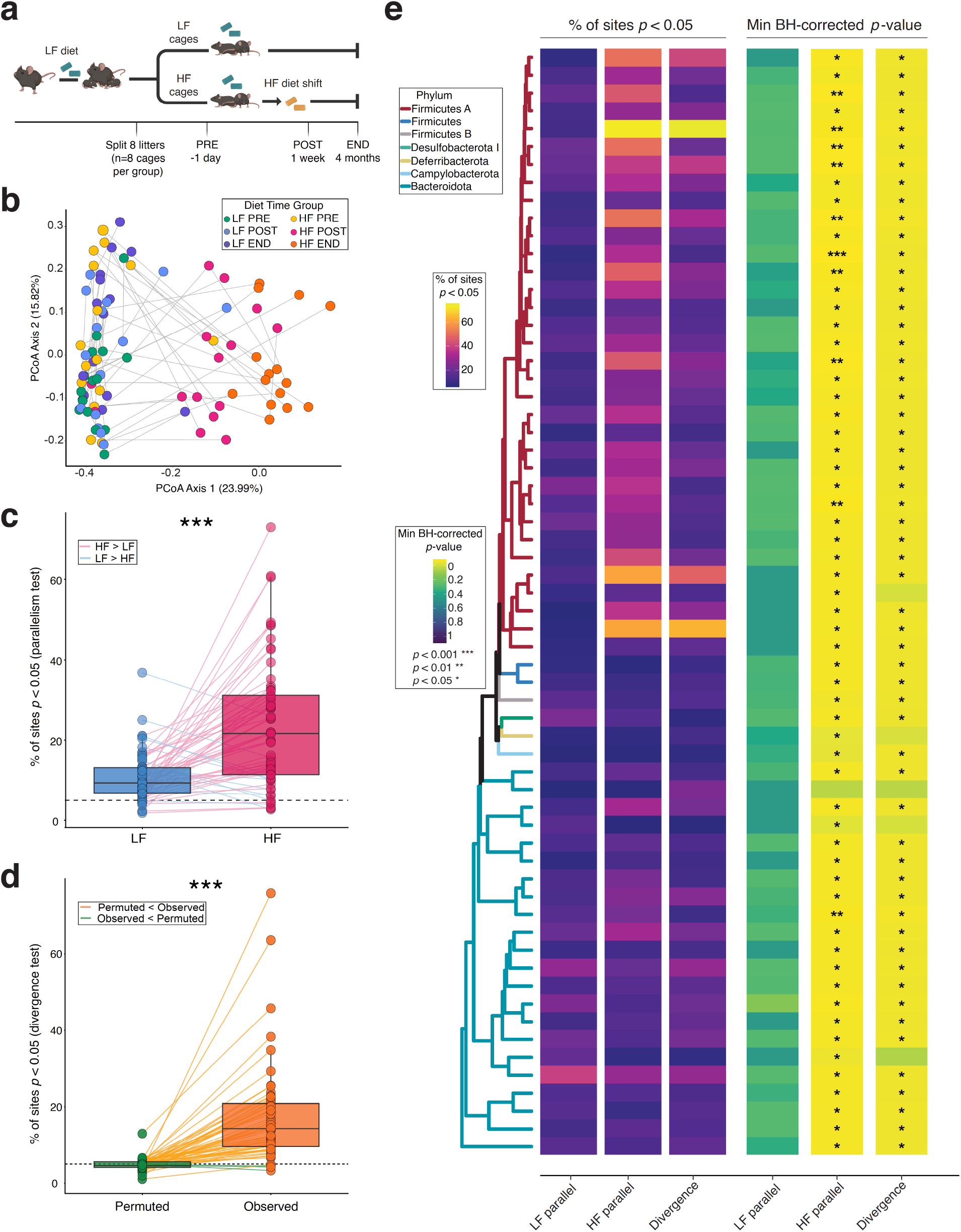
Widespread adaptive evolution of gut bacterial species in response to high-fat/low-carbohydrate diets. **a**, Cartoon shows overview of experimental design, in which litter-matched mice were transitioned to a high-fat/low-carbohydrate (HF) diet or maintained on their ancestral low-fat/high-carbohydrate (LF) diet. **b,** Principal coordinates analysis plot shows the Bray-Curtis dissimilarities of species composition among metagenomes from HF- and LF-fed mice at the three deeply sequenced timepoints (PRE, POST, and END). Each point represents a sample, and points are colored by treatment group and timepoint, as indicated by the key. Lines connect samples from the same mouse. **c,** Boxplots show the percentage of variable sites in bacterial species displaying parallel allele frequency shifts across replicates, based on a *p* < 0.05 threshold (t-test for directional mean allele frequency shift), in LF or HF groups. Points represent species, and lines connect the same species between LF and HF groups. Colors of lines denote the direction of difference between LF and HF, as indicated by the key. **d**, Boxplots show the percentage of variable sites in bacterial species displaying divergent allele frequency shifts between LF and HF groups, based on a *p* < 0.05 threshold (paired t-test for differences in mean allele frequency shift). Pairs of points represent species, and lines connect the same species between the real dataset (observed) and a permuted dataset in which sample-to-timepoint metadata assignments within each diet group were randomized. Colors of lines denote the direction of difference between observed and permuted datasets, as indicated by the key. In **c** and **d**, asterisks indicate significance of differences in means between boxplots; One-sample t-test *** *p* < 0.001. **e**, Phylogeny shows relationships among 62 core gut bacterial species tested for parallel and divergent allele frequency changes mediated by HF diets. Colors of branches denote phylum assignments. Leftmost three columns of heatmap show, for each species, the % of sites with *p* < 0.05 in LF parallelism, HF parallelism, or divergence tests. The three rightmost columns of heatmap show, for each species, the minimum BH-corrected *p*-value of all tested sites in that species’ genome derived from LF parallelism, HF parallelism, or divergence tests. Asterisks indicate minimum BH-corrected *p*-values; * *q* < 0.05, ** *q* < 0.01; *** *q* < 0.001.

## Widespread reproducible adaptive evolution of gut bacterial species across replicates

We leveraged the genome-resolved metagenomic dataset to test for reproducible adaptive evolution of gut bacterial species across experimental replicates (i.e., mouse litters) in response to the transition to a HF diet. First, we created a software toolkit for community-wide allele-level population genetic analyses (https://github.com/MoellerLabPU/AlleleFlux) and used it to profile allele frequencies at every genomic position in all 160 species in every mouse at every sampling timepoint. We applied conservative filters to mitigate effects of sequencing errors and erroneous mapping that may bias allele-frequency estimates (see Methods). We identified 62 species present in each diet group in all eight replicates at both the PRE and END timepoints. The genomes of these core species contained 482,336 variable sites, which we defined as genomic positions whose nucleotide identity varied within mice between PRE and END timepoints in at least 50% of replicates in at least one diet group (HF or LF).

Using these data, we tested for parallel evolution—a classic hallmark of deterministic adaptation ^24,36^—of gut bacterial species across replicates in response to HF diets. We calculated the mean change in frequency of every nucleotide at every variable site in each of the 62 core species between the PRE and END samples for each cage of mice (see Methods). Under genetic drift, the expected change in an allele’s frequency is zero. In contrast, positive or negative selection would be expected to drive consistently positive or negative changes in allele frequency across mouse cages (i.e., parallel changes across hosts).

We first tested whether the mean change in each allele’s frequency within cages differed significantly from zero within each diet group (LF or HF) (two-sided t-test *p* < 0.05). Under the null hypothesis and assuming independence among sites, 5% of sites in each species were expected to display *p* < 0.05. However, due to linkage among sites within a single genome, not all sites are independent, such that selection at one site may drive parallel allele-frequency changes not only at that site but also at linked sites via ‘genetic hitchhiking’, whereby linked alleles change in frequency together due to selection on a subset of the alleles. In line with this possibility, ∼90% of core species (56/62) displayed parallelism at > 5% of variable sites in the HF group at *p* < 0.05 (Fig. 1c; Supplementary Table 2; Supplementary Data 1), with some species showing parallel changes in allele frequency at over half of all variable sites based on a *p* < 0.05 threshold (Fig. 1c; Supplementary Table 2; Supplementary Data 1). The extent of parallelism across species was significantly higher in the HF group compared to the LF group (Fig. 1c; paired t-test, HF score > LF score, *p* = 5.26e-8; Supplementary Data 1). Next, we conducted false-discovery rate corrections using the Benjamini–Hochberg (BH) method across all tested sites. These analyses allowed us to ask whether each species’ genome showed any significant evidence of parallel allele frequency changes after correcting *p*-values for multiple tests. Remarkably, of the 62 core species, 61 contained at least one variable site whose allele frequencies displayed significant parallelism across replicates in response to the HF diet after accounting for multiple testing (*q* < 0.05; Fig. 1e). In contrast, none of these species displayed significant parallelism in response to LF diets after accounting for multiple testing. Given widespread significant evidence of parallel evolution of gut bacterial species across replicates in response to HF diets, we next tested whether species displayed divergent allele frequency changes between diet groups. To address this question, we tested whether mean within-cage allele frequency changes between END and PRE timepoints differed significantly between diet groups. We found that ∼94% (58/62) of species displayed divergence in allele frequencies at > 5% of tested variable sites at *p* < 0.05 (t-test, paired by replicate, *p* < 0.05; Fig. 1d; Supplementary Table 2; Supplementary Data 1; see Methods). The extent of allele-frequency divergence observed in the real data significantly exceeded that observed when analyses were conducted after permuting sample-to-timepoint metadata assignments within each diet group (paired t-test, observed > permuted, *p* = 2.2e-16; Fig. 1d; Supplementary Table 2; Supplementary Results). After BH correction for multiple testing, we observed significant evidence of divergent allele frequency trajectories between diet groups in at least one variable site in 57 of the 62 core species (*q* < 0.05; Fig. 1e). The observation that 61/62 and 57/62 species showed significant evidence of parallelism and divergence (based on a minimum per-site *q*-value < 0.05), respectively, was robust to the exclusion of sites with mean sequencing coverage greater than two standard deviations from its species’ mean coverage and sites within mobilome genes (see Methods; Supplementary Data 1), exclusions designed to minimize mismapping artefacts.

Alleles showing both significant BH-corrected divergence between diet groups and significant BH-corrected parallelism within the HF group predominantly represented standing genetic variation detected in mice at the PRE timepoint, rather than putative *de novo* mutations. Across the 57 species containing at least one allele significant in both tests, the median number of these significant sites was 120 per species (Supplementary Data 1). Only 20 of these species contained any significant alleles that were not detected at the PRE timepoint and were therefore classified as putatively *de novo*. Among these 20 species, the median number of putative *de novo* mutations was five (∼ 4.2% of the median number of significant sites per species). The median across species of their mean one-read detection limit was 0.251%, calculated as 1/N, where N is the number of high-quality reads mapped to the position at the PRE timepoint pooled across mice. Moreover, for each of the 61 species showing significant evidence of HF-mediated parallelism (Fig. 1), each had an allele detected at the PRE timepoint that showed BH-corrected significant evidence of parallelism. Thus, the reproducible allele-frequency responses to the HF diet transition arose primarily from standing genetic variation.

To further evaluate the robustness of our results, we conducted parallelism and divergence tests using several other well-established methods for detecting adaptive evolution in E&R experiments (see Methods) including Wilcoxon tests, Cochran–Mantel–Haenszel tests, and linear mixed effects models. These analyses confirmed widespread parallel and divergent allele-frequency shifts in response to HF diets (Extended Data Figs. 6–8; Supplementary Table 2; Supplementary Results) and suggested that t-tests (Fig. 1) were most conservative (least powerful and least prone to false positives) of the parametric methods. Widespread significant parallelism and divergence of allele-frequency changes were also observed when analyses shown in Figure 1 were conducted using a more stringent threshold of a minimum mean coverage of 1x in all cages at each PRE, POST, and END timepoint (Extended Data Fig. 9). In addition, we observed no significant relationships between species’ relative abundances (as estimated by coverage in inStrain^41^) or change in relative abundances (END vs PRE) and the minimum *p*- or *q-*values obtained from HF parallelism or HF–LF divergence analyses for each species (Spearman rank correlation *p*-value > 0.05 for all tests).

Cumulatively, these results indicate that transitions to HF diets drove reproducible within-host adaptive evolution of essentially all core gut bacterial populations. Besides selection, the only other of the four evolutionary forces (selection, mutation, migration, and genetic drift) capable of producing directionally consistent changes in allele frequencies across replicates are migration and mutation. However, in our experiment, migration—in this context the transmission of microbiota between cages or from the external environment—was held constant by rearing HF and LF cages on their respective sterile diets in the same mouse facility. Similarly, differences in mutation rate mediated by HF diets cannot readily explain our results, because alleles showing significant parallelism in the HF group and divergence between HF and LF groups largely represented standing variants present at the PRE timepoint. Thus, the observation of parallel evolution across HF cages (but not across LF cages) and divergent evolution between HF and LF cages is indicative of deterministic evolutionary responses of bacterial populations to selection, *i.e.*, adaptive evolution. Sites within the same genome may evolve in linkage disequilibrium, such that many of the sites detected may not be under selection themselves but instead linked to a site under selection. However, the detection of at least one site showing BH-corrected significant HF parallelism or HF–LF divergence in nearly every (61/62) core gut bacterial species is indicative of widespread responses to selection mediated by the HF dietary transition.

## HF diets reproducibly drove strain replacements within bacterial species

That some species showed genome-wide allele-frequency changes in response to HF diets (Fig. 1c and d) is indicative of genetic hitchhiking. In the extreme, these dynamics can arise due to strain replacement or strain sorting mediated by the HF diet. In this mode of adaptive evolution from standing variation, multiple strains of a species inhabited individual mice at the beginning of the experiment, and a subset of those strains were positively selected following the HF dietary transition.

We tested for adaptive strain sorting in two ways. First, we compared the END and PRE samples from each cage of mice and tested for each of the 62 species shown in Fig. 1 whether the species experienced a change in the dominant strain within each cage, as defined by a mean within-mouse consensus ANI (conANI) of less than 99.9% (the strain-level conANI threshold recommended by inStrain^41^) (Supplementary Table 5). We then used a permutation Chi-squared test with BH correction for multiple testing across species to identify species that displayed more frequent changes in the dominant strain in either HF cages or LF cages. These analyses revealed a single species, a Lachnospiraceae (UBA3282 sp009774655), that showed significantly more frequent strain turnover in HF cages than in LF cages (BH-corrected Chi-squared *p* = 0.0073) (Fig. 2). No species showed significantly more frequent strain turnover in LF cages than in HF cages. Second, we tested for each bacterial species whether populations of strains from HF-fed mice from different cages were more genomically similar at the END timepoint than expected based on the mean population ANI (popANI) between HF and LF cages. These tests effectively asked whether genomic similarities among populations of strains clustered by HF diet. After BH correction, we found that ∼48% of species displayed significant evidence of HF-mediated genomic clustering of populations across replicates (Supplementary Table 5; Supplementary Results). Again, Lachnospiraceae sp009774655 displayed the strongest evidence of HF-mediated genomic clustering of populations (BH-corrected permutation t-test *p* < 0.05; Fig. 2; Supplementary Results). Incorporating data from shallow sequencing on fecal samples collected every ∼2 weeks (Extended Data Fig. 1; Methods) revealed that alleles at different variable sites of this species displayed coordinated shifts in frequency over time (Fig. 2d, e), indicating linkage. The selection on this strain mediated by the transition to HF diets represents an example of adaptive evolution from standing variation, as the alleles that rose to high frequency in the HF group by the END timepoint were present at non-zero frequency at the PRE timepoint in both diet groups (Fig. 2d, e). This species was also the only species that contained genes that showed significantly elevated ratios of non-synonymous to synonymous substitution rates (i.e., dN/dS > 1) across replicates in the HF-diet group based on comparisons between END and PRE timepoints (Fig. 2c; see Methods). These genes coded for glycosyltransferase, which catalyzes glycosidic linkages, and Appr-1″-pase, which regulates tRNA splicing. These results are consistent with the possibility that positive selection predating our experiment contributed to historical gene divergence between the strains that displayed differential fitness in response to HF and LF diets. Cumulatively, these findings indicate that HF diets reproducibly drove adaptive strain replacements in gut bacterial species within individual mice.

**Fig. 2:**
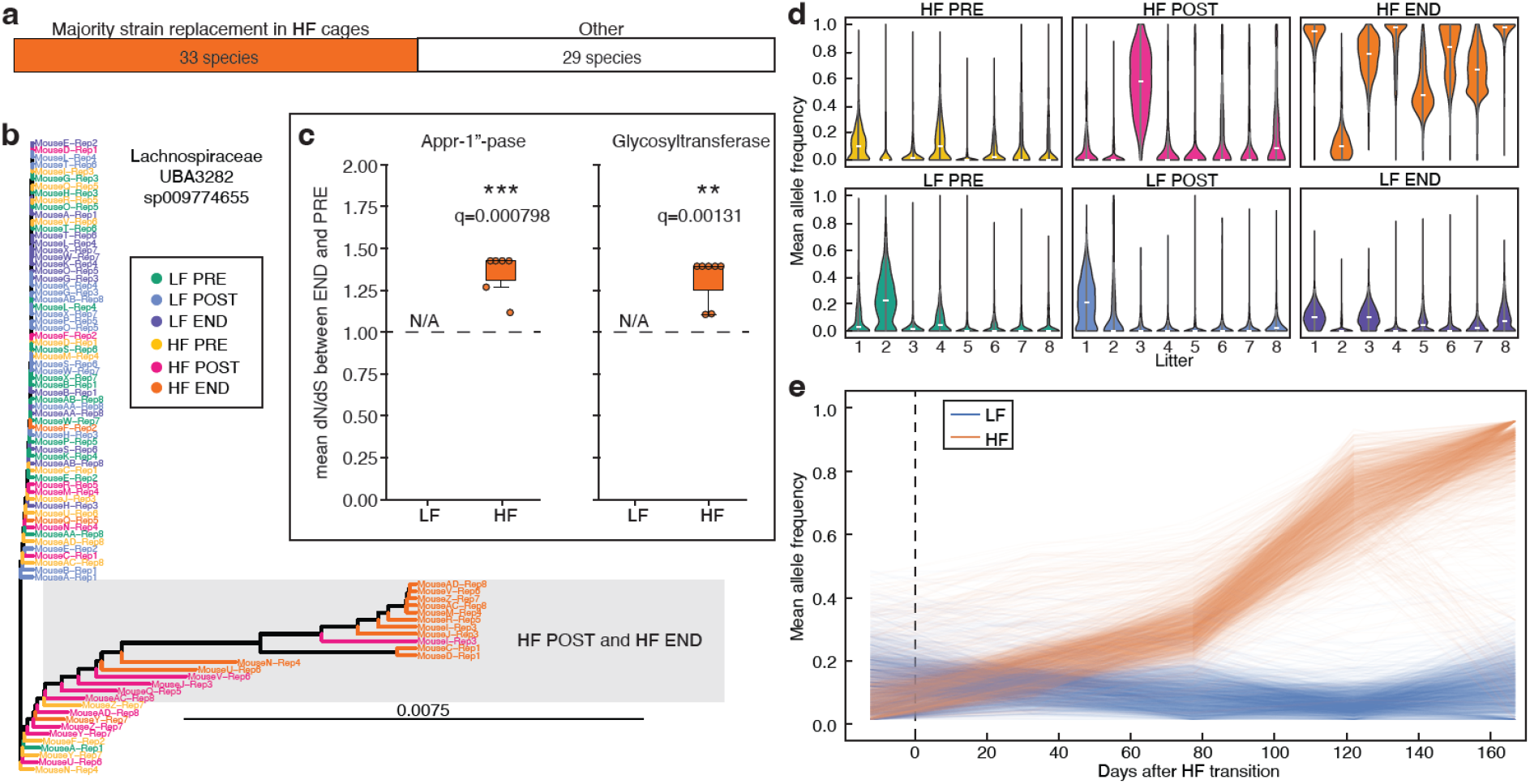
High-fat/low-carbohydrate diets drove strain-replacement events within bacterial species. **a**, Stacked bar chart shows number of species displaying evidence of changes in the dominant strain between END and PRE timepoints in a majority of HF replicates, based on a conANI threshold of 99.9%. **b**, Dendrogram shows genomic similarity (conANI) among the dominant strains of Lachnospiraceae UBA3282 sp009774655, the species which showed the strongest evidence of diet-mediated strain replacement, across samples. Each tip represents sp009774655 in a metagenomic sample, and tip labels and terminal branches are colored by treatment group and timepoint, as indicated by the key. **c**, Boxplots show mean per-replicate dN/dS in LF- and HF-fed mice of the two genes that showed significantly elevated (> 1) dN/dS across replicates in sp009774655. Each point represents the mean of dN/dS for a replicate, as calculated by comparing the END and PRE timepoints for each mouse within the replicate. N/As indicate no changes in sequence within LF-fed mice. Asterisks indicate significance of BH-corrected *p*-values (t-test for mean > 1); ** *q* < 0.01, *** *q* < 0.001. **d**, Violin plots show for sp009774655 within each litter at PRE, POST, or END time points the allele frequencies of major alleles detected in the HF END timepoint. Widths of violins indicate density and horizontal bars within violins indicate medians. **e**, Line plot shows allele frequency trajectories of variable sites in sp009774655 showing BH-corrected significant evidence of allele-frequency divergence between HF and LF groups. Each line indicates the mean allele frequency across replicates of the allele showing the greatest mean positive change in frequency in HF cages between PRE and END timepoints.

## Genomic responses to HF diets were enriched in a subset of gene families across species

We next tested whether any gene functions were enriched for signatures of deterministic responses to HF diets across gut bacterial species, as expected if species shared genomic bases of adaptation in response to HF diets. To investigate gene-wise adaptive responses to HF diets, we focused these analyses only on the 29 species that did not show strain replacement in >50% of replicate HF cages (Supplementary Table 6), where strain replacement was defined as a mean within-mouse consensus ANI (conANI) of less than 99.9%. Within these species, we identified every gene that contained a site showing significant allele-frequency divergence between HF- and LF-diet groups and parallelism in the HF-diet group based on BH-corrected *p*-values. In total, these species contained a median of 14 of these significant sites per species representing a median of 4 contiguous sequences (contigs) per species (Supplementary Table 6). We then conducted hypergeometric tests to identify annotations (Clusters of Orthologous Groups, COG20) that were significantly enriched among this set of genes relative to the complete set of these species’ genes tested (Supplementary Table 7). These analyses allowed us to test for parallel responses to selection across different species within hosts. Results revealed multiple significantly enriched functional categories and protein descriptions after BH correction across annotations (Fig. 3a). Enriched functional categories included: Replication, recombination and repair; Transcription; Cytoskeleton; and Signal transduction mechanisms. Significantly enriched protein descriptions included multiple recombinases, invertases (Fig. 3b, c), and others. Examples of the helix-turn-helix protein family enriched among the set of genes showing allele-frequency divergence are shown in Fig. 3d. Together, these results indicate that HF diets drove deterministic allele-frequency changes in a common set of genomic functions and pathways across distantly related gut bacterial species.

**Fig. 3:**
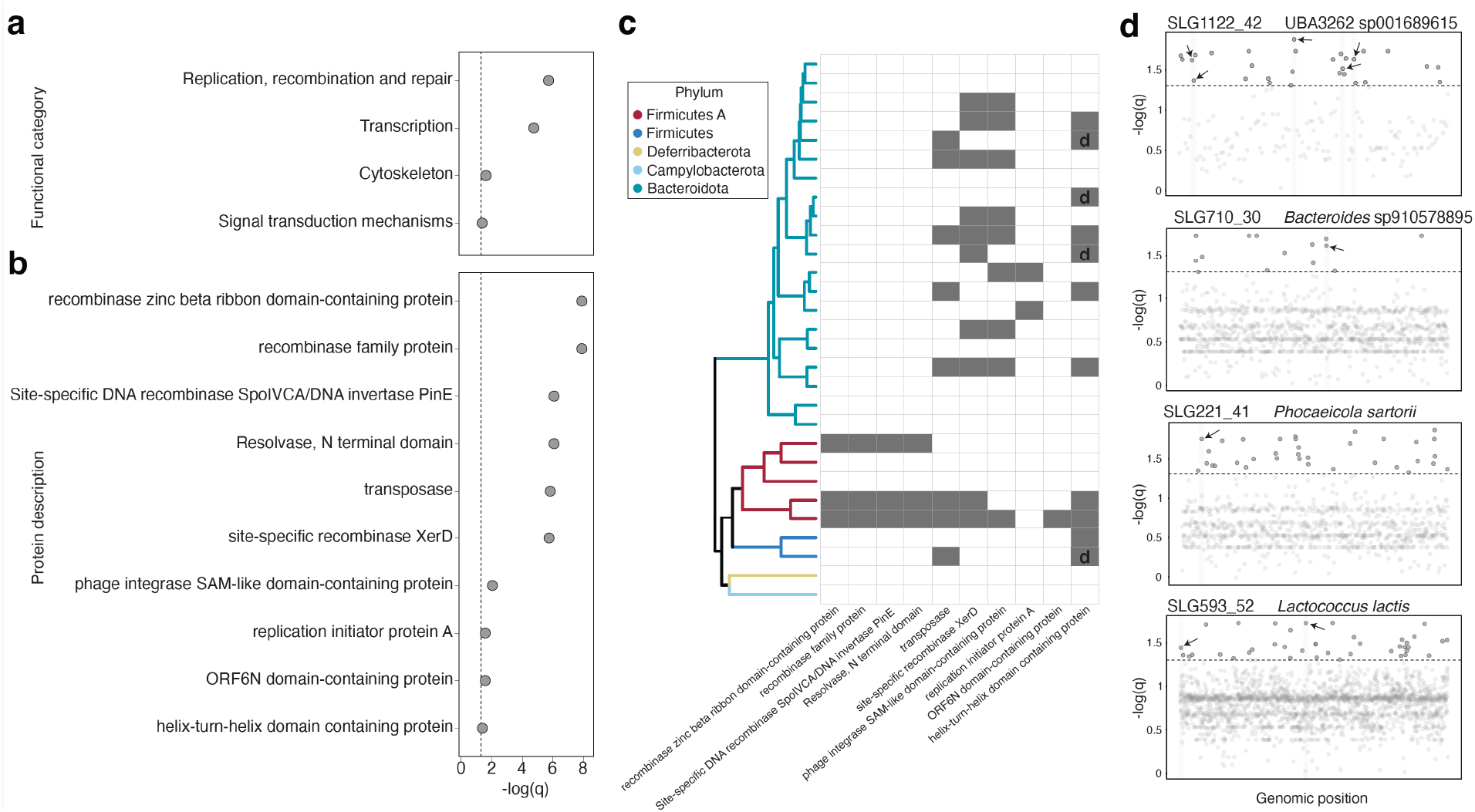
Parallel signatures of positive selection across bacterial species. **a** and **b,** Plots show the strength of significance (-log of q-value) for enrichment of specific functional categories and protein annotations among the set of genes containing sites showing BH-corrected significant allele-frequency divergence between HF and LF groups and parallelism in the HF group (compared to the total set of genes tested). **c**, Phylogeny shows relationships among species that did not show strain replacement in a majority of HF cages between END and PRE based on a conANI threshold of 99.9%. Grid right of phylogeny denotes the per-species presence (grey) or absence (white) of genes showing significant parallelism in the HF group and divergence between HF and LF group based on BH-corrected *p*-values. Cells in the last column labeled ‘d’ correspond to Manhattan plots **(d). d**, Manhattan plots show degree of significance of allele-frequency divergence at each variable site in four representative gut bacterial species that showed parallel significant allele-frequency divergence in the helix-turn-helix domain containing protein gene family (arrows). MAG IDs and species names are shown above each plot. Dashed horizontal lines indicate significance cutoff at BH-corrected *p*-value < 0.05.

## A subset of alleles rose in frequency in HF mice after declines in population diversity

Given that the transition to the HF diet led to selection on bacterial populations, as evidenced by reproducible allele-frequency changes across experimental replicates, we next investigated whether any populations experienced changes in overall levels of genetic diversity following the HF dietary transition. To test this idea, we calculated nucleotide diversity (π), which measures the degree of genetic differences between randomly drawn sequences from a population in a sample, for each bacterial species in each sample from PRE, POST, and END timepoints using inStrain^41^ and a rarefaction depth of 50x coverage per site. We used two-sided Welch t-tests for differences in π between HF and LF cages within each species and timepoint (PRE, POST, or END), then applied BH correction of p-values across species within each timepoint. These analyses revealed that 0/62 species showed significant differences (Welch t-test; *q* < 0.05) between HF and LF in the PRE timepoint (as expected), 2/62 showed significant differences at the POST timepoint, and 10/62 showed significant differences at the END timepoint (Supplementary Table 8; Extended Data Fig. 10). The only two species that showed significant differences in π at the POST timepoint, *Odoribacter* sp910578105 and *Cryptobacteroides* sp009774765, each showed significant reductions in π in the HF group at the POST timepoint, did not show evidence of strain turnover in any replicate (based on a conANI threshold of 99.9%), and had only 2 and 3 variable sites (each on different contigs), respectively, showing evidence of both HF parallelism and HF–LF divergence in allele frequency trajectories. For both *Cryptobacteroides* sp009774765 and *Odoribacter* sp910578105, these alleles were detected at low frequency in mice at the PRE timepoint, indicating selection on standing variation. In *Cryptobacteroides* sp009774765, these sites were in an intergenic region, a gene annotated as an ATPase, and a gene annotated as a hexosaminidase/glycosyl hydrolase (Supplementary Table 6; Supplementary Data 1). The latter annotated protein description is suggestive of functions related to the breakdown of complex carbohydrates and/or glycolipids. These sites in *Odoribacter* sp910578105 were in two genes, one annotated as a phage integrase and the other as an Arm DNA-binding domain (Supplementary Table 6; Supplementary Data 1). These results are consistent with purging of genetic diversity in these species after the HF dietary transition, leading to reductions in π, and positive selection mediated by the HF dietary transition acting on a small number of standing polymorphisms.

## HF diets reproducibly altered phase variation within invertons

Given that allelic variation within genes coding for recombination machinery responded to the HF dietary transition (Fig. 3), we investigated HF diet–induced changes in structural variation within bacterial genomes. For these analyses, we cultivated, isolated and sequenced bacterial strains from fecal samples collected from HF and LF groups at PRE, POST, and END timepoints. Relative to MAG-based analyses, which could be sensitive to mismapping and assembly artefacts, analyses based on isolate genomes enabled more robust detection of structural variation^43^. We employed culture conditions to recover gut Bacteroidales^43^, whose genomes harbor environmentally induced structural variation. For example, *Bacteroides* genomes contain invertons—DNA segments whose orientation can invert reversibly—that respond adaptively to environmental cues^42,44,45,46,47,48,49^. Invertons are flanked by inverted repeat sequences targeted by site-specific recombinases and can act as switches that alter the expression of nearby genes or create new coding regions. Despite their potential importance for bacterial adaptive evolution, the diet-induced responses of Bacteroidales invertons within complex gut microbiota are poorly understood.

Culturing, isolation and whole-genome sequencing of 562 isolates yielded 79 assemblies from *Phocaeicola sartorii* and 43 assemblies from *Bacteroides muris* (Supplementary Table 9), the two species that we recovered from PRE and END timepoints from HF- and LF-diet groups. The genomes of these isolates confirmed the adaptive allele-frequency responses to HF dietary transitions detected in metagenomic data (Supplementary Results). Using these genomes, we identified invertons in *P. sartorii* and *B. muris* and assessed their responses to HF diets. Using PhaseFinder^45^, we identified 82 *P. sartorii* invertons and 68 *B. muris* invertons that displayed variation in the orientation of the invertible region *in vitro* (Supplementary Table 10, see Methods). Using metagenomic data generated from fecal samples, we applied linear mixed effects models and BH correction across invertons to test whether these *P. sartorii* and *B. muris* invertons significantly changed orientation within mouse metagenomes in response to HF diets (Supplementary Table 11). These analyses revealed two invertons whose orientation significantly changed between PRE and END timepoints in one diet group but not the other. Both invertons were within 100 kilobases in the *P. sartorii* genome and significantly reversed orientation across replicates after the transition to the HF diet but did not change orientation in the LF-diet group (Fig. 4a). Importantly, zero HF cages exhibited evidence of a change in the dominant strain within *P. sartorii* between END and PRE timepoints (mean within-mouse END–PRE conANI > 99.9% for all HF cages), indicating that changes in the orientation of invertons occurred without detectable replacement of the dominant strain under our conANI criterion. Inspection of allele frequencies near the extremes of these invertons confirmed diet-induced changes in orientation (Supplementary Results). To assess functions of these invertons, we examined the promoters^50^ and genes within and flanking the invertible region. For the inverton spanning sites 108998–109206, the HF diet–mediated inversion swapped a predicted weaker promoter for a predicted stronger promoter upstream of a cluster of five genes of unknown function (Supplementary Results). For the inverton spanning sites 209006–209503, the HF diet–mediated inversion swapped a predicted weaker promoter for a predicted stronger promoter upstream of the glycan foraging genes *susC* and *susD* (Supplementary Results). Thus, the dietary transitions altered predicted regulatory regions of known carbohydrate foraging genes in *P. sartorii*. Specifically, the inversion localized a predicted promoter sequence upstream of carbohydrate foraging genes *susC* and *susD* under relatively low-carbohydrate (HF) conditions. Cumulatively, these results demonstrate invertible phase variation reproducibly responds to changes in host diet within endogenous mammalian gut microbiota.

**Fig. 4:**
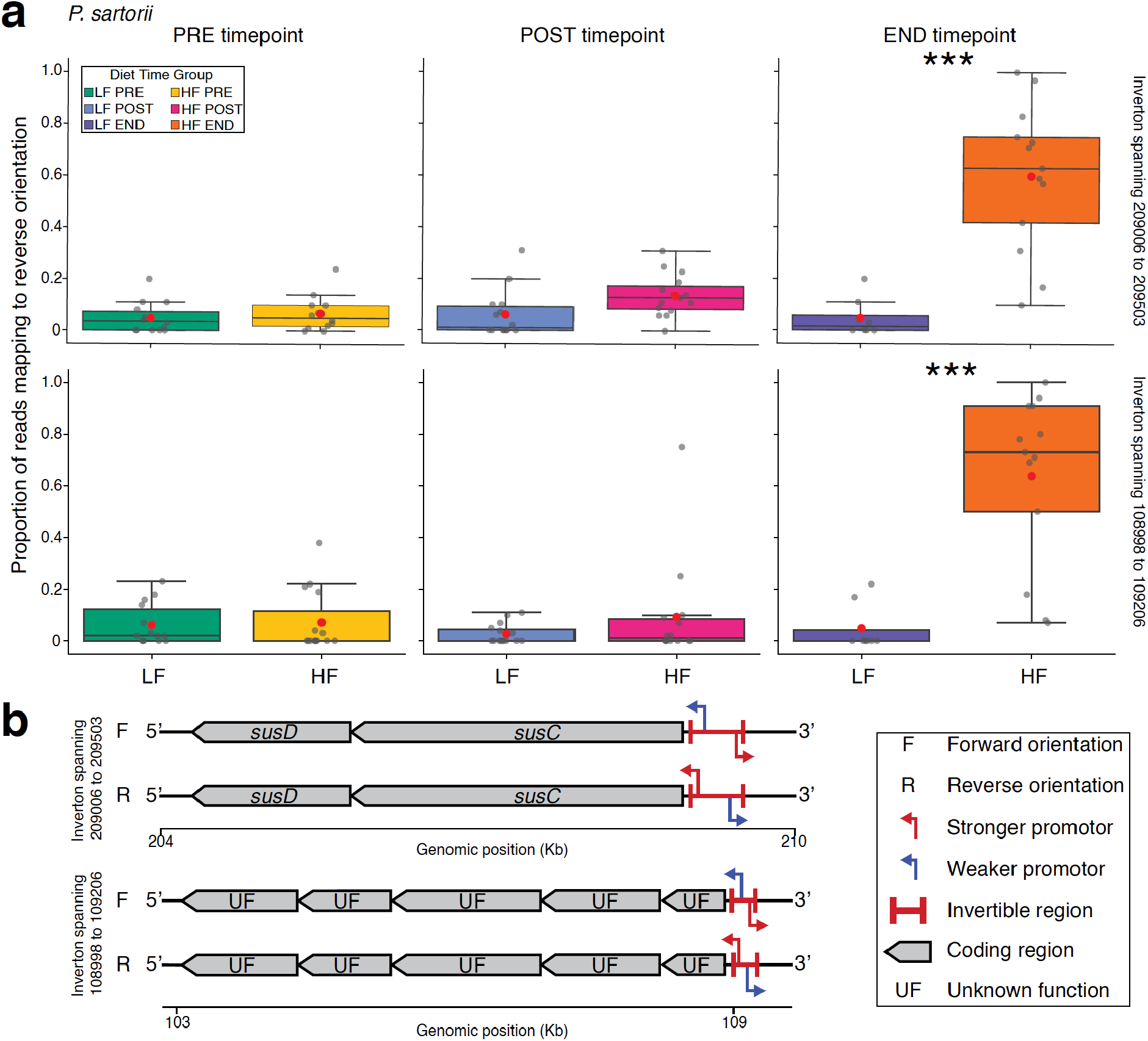
Invertons in *P. sartorii* respond reproducibly to high-fat/low-carbohydrate diets. **a**, Boxplots show proportion of inverton-derived sequences displaying the reverse orientation in metagenomic data from LF- and HF-fed mice at PRE, POST, and END timepoints. Each row of plots corresponds to an inverton. Each point represents a mouse. Red points denote means of boxplots. Colors of boxplots correspond to diet group and timepoint, as indicated by the key. Asterisks indicate BH-corrected significant differences between HF and LF groups based on linear mixed effects models controlling for replicate (i.e., litter); *** *q* < 0.001. **b**, Cartoons show nearby genes and promoters within the invertible region of invertons from **a**. Cartoons depict relative positioning of invertible regions, genes, and promoters. ‘F’ and ‘R’ denote forward and reverse orientations of the invertible region; red and blue bent arrows denote predicted ‘stronger’ and ‘weaker’ promoters, respectively, as identified by BPROM^50^; bold red lines denote invertible regions; grey boxes denote coding regions; and ‘UF’ denotes unknown function, as indicated by the key.

## Conclusions

Investigating population genomic responses of complex gut microbiota within an isogenic, native host genetic background allowed us to quantify adaptive population-genetic responses of gut bacterial populations to dietary transitions within individual hosts. The observed diet-mediated deterministic evolution cannot be explained by the non-selective evolutionary forces (mutation, migration, or genetic drift). Histories of migration were controlled by litter-matching HF and LF mice and maintaining them in the same animal facility. Likewise, our results do not appear to reflect changes in mutation rates imposed by HF diets: nearly all alleles showing deterministic allele frequency changes were present in mice before the dietary transition (Supplementary Data 1). Changes in genetic drift cannot explain reproducible changes in strain or allele frequencies across replicates. Thus, our results indicate reproducible community-wide adaptive responses of gut bacterial populations to the HF dietary transition.

In principle, adaptive responses could confer fitness advantages at one or more levels of biological replicators, including genes, mobile elements, or bacterial lineages. In our experiment, responses to selection included both strain sorting and gene-specific allele-frequency changes. In multiple species, HF diet–mediated selection drove differential fitness and sorting of standing strain diversity, as evidenced by genome-wide changes in allele frequencies in parallel across experimental replicates (Figs. 1 and 2). In species that did not exhibit clear strain sorting or replacement, signatures of HF diet–mediated selection were enriched in specific functional categories and protein families (Fig. 3), indicating parallel responses to selection across gut bacterial species within hosts. A subset of gene-specific allele-frequency changes could be attributed to inducible structural variation, which we demonstrated by combining analyses of genome-resolved metagenomes and isolate genomes (Fig. 4). The adaptive responses of gut bacterial species may be due to direct selection imposed by the dietary transition, but they may also reflect indirect effects mediated by the dietary transition (e.g., changes to interactions between bacterial lineages). Cumulatively, our study establishes the community-wide scope and reproducibility of diet-mediated adaptive evolution in the gut microbiota.

## Supporting information

Supplementary Information Guide

Supplementary Results

## Methods

### Animal experiments

All mouse experiments were approved by Cornell University Institutional Animal Care and Use Committee Protocol #2015-0060. To enable tests for reproducible microbiota evolution in response to transitions to high-fat/low-carbohydrate (HF) diets, we conducted E&R experiments using replicated mouse litters derived from a single pair of grandparents. The grandparental breeding pair was selected from a wild-derived inbred line (NY3) of *Mus musculus domesticus* mice and maintained *ad libitum* on a standard lab chow diet (TEKLAD Irradiated LM-485 Mouse/Rat Diet, kcal from macronutrients: 17% fat, 25% protein, 58% carbohydrates). This breeding pair yielded four breeding pairs, each composed of littermates. These breeding pairs were transitioned after weaning to a purified low-fat/high-carbohydrate (LF) diet (Envigo Teklad Custom Diet TD.120582) designed to match the macronutrient breakdown of the chow diet. When these four breeding pairs had pups, four mice from each litter were weaned into two cages, with two mice per cage and each cage containing the same sex distribution (e.g., male-female, male-male, or female-female). Thus, all mice were matched by sex within each experimental replicate (i.e., mouse litters). For each replicate (i.e., each of eight litters for which sampling was conducted as shown in Fig. 1a), both cages of littermate mice were weaned onto the LF diet. Then, after one week, one cage selected at random was switched to the purified HF diet (Envigo Teklad Custom Diet TD.210072). The HF diet was identical to the LF diet in micronutrient breakdown, but the HF diet had > 4x the number of kilocalories derived from fat (Extended Data Table 1). The LF diet had the following kcal from macronutrients profile: 17.2% fat, 18.8% protein, 63.9% carbohydrates. The HF diet had the following kcal from macronutrients profile: 74.4% fat, 13.2% protein, 12.4% carbohydrates. Food and water were provided *ad libitum*. Mice were maintained on these diets for ∼4 months.

### Sample collection and metagenomic sequencing

For fecal collection, mice were temporarily moved into individual holding cages, and fresh fecal samples were collected from the bottom of holding cages within 20 minutes. All samples were immediately frozen on dry ice. DNA was extracted from all samples using a bead beating protocol (Qiagen PowerLyzer PowerSoil Pro), and libraries were prepared with a TruSeq-equivalent library preparation at the Cornell Biotechnology Resource Center or Hackflex library preparation, as previously described^23,51,52^. TruSeq and Hackflex libraries were each sequenced on one lane of Illumina NovaSeq 6000 at the UC Davis Genome Center.

### Genome assembly and database creation

Raw reads were assembled into metagenome-assembled genomes (MAGs) with a custom snakemake pipeline (github.com/CUMoellerLab/sn-mg-pipeline) as previously described^23^. Briefly, adapter sequences were removed with cutadapt^53^ (v0.17.4), host reads were removed by mapping to the mouse refseq genome (GCF_000001635.27/) with bowtie2^54^ (v2.3.0), and quality of sequences was assessed with fastqc^55^ (v0.72.0). Sequences were assembled into contiguous sequences (contigs) with MEGAHIT^56^ (v1.2.9). Before binning contigs into MAGs, quality-filtered reads were mapped against each set of contigs derived from samples collected from mice of the same litter. This mapping strategy was chosen because littermates were expected to share strains. All-by-all mapping of reads to contigs among samples from littermates was performed with minimap2^57^ (v2.24). Using the coverage information obtained from mapping, contigs were then binned with three binners: metaBAT 2^58^ (v2.15), MaxBin 2.0^59^ (v2.2), and CONCOCT^60^ (v1). The optimal set of bins for each sample was then selected with DASTool^61^ (v1.1.3), yielding a total of 4,313 MAGs. 843 chimeric MAGs were removed with GUNC^62^ (v1.0.6). MAGs estimated to be > 90% complete and < 5% contaminated by CheckM2^63^ (v1.0.1) were identified, yielding 2,218 high-quality MAGs, which were used for downstream analyses.

To identify species, MAGs were dereplicated based on 95% shared average nucleotide identity (ANI) with dRep^64^ (v3.4.2). The genomes dereplicated into 160 unique species genome clusters (“species”), each typified with a representative genome. These representative genomes formed the reference genome database that was used for downstream analyses. A phylogeny of these MAGs (Extended Data Fig. 2) was built using IQ-TREE^65^ (v2.2.2.6) using the GTDB-Tk^66^ (database release v207 v2) alignment with model selection restricted to “WAG, LG” and 1000 bootstrap replicates. The model “LG+R10” was selected. In addition, MAGs were classified to taxonomic ranks with GTDB-Tk^66^ (v2.3.2, database release 207 v2) using the “align”, “classify”, and “identify” commands.

### Diversity and abundance analyses

The relative abundance of each reference genome in each sample was calculated using results of mapping analyses from bowtie2^54^. The total number of reads in each sample was determined with “seqkit stats” from seqkit^67^ (v0.15.0). The reads mapped to each species were determined via the “filtered_read_pair_count” as reported by “profile” run in “database mode” in inStrain^41^. Relative abundances for each species were normalized by genome coverage within each sample. Species not detected in a sample were assigned relative abundance values of 0 for that sample.

All alpha and beta diversity metrics and statistics were calculated in R with vegan^68^ (v2.6.8). Samples were rarefied to 6,832,952 reads (the minimum sequencing depth across all samples) with “rrarefy” and read counts per species were normalized to relative abundance within each sample by dividing by total quality-filtered read pairs and genome size, then rescaling to sum to 100, prior to calculating alpha and beta diversity metrics. Taxa barplots (unrarefied data) were created and visualized using ggplot2^69^ (v3.5.1) (Extended Data Fig. 3). Shannon diversity (Extended Data Fig. 5) was calculated with the “diversity” function, and medians were compared with Wilcoxon tests and one-tailed Wilcoxon tests. Bray-Curtis and Jaccard dissimilarities were calculated with “vegdist” and PERMANOVA was performed with “adonis2”. PCoAs were generated with the “cmdscale” function from the base R “stats” package (v4.4.1) and were visualized with ggplot2. Differential species-abundance testing was performed using the ANCOM-BC2 function from the ancombc package^70^ (v2.6.0) with default settings except: fix_formula = “diet_time”, random formula = “(1|cage) + (1|sex)”, and dunnet = “TRUE”, where “diet_time” corresponded to the group (HF or LF) and timepoint (PRE, POST, END) for each sample.

### Allele frequency profiling of species

We created a custom pipeline for profiling allele frequency changes. (https://github.com/MoellerLabPU/AlleleFlux). We quantified allele frequencies for each sample at each position in reference genomes of species detected in the sample. Species genome clusters were counted as present in a sample if the reads from that sample yielded breadth > 10%. Per-sample allele frequency profiles were generated from sorted BAM files aligned to a reference FASTA containing contigs of all species’ representative MAG (95% ANI clusters, identified by dRep^64^). For each genomic position with non-zero coverage, pysam’s (https://github.com/pysam-developers/pysam) pileup engine (stepper=“samtools”) was used to record aligned bases, individual nucleotide counts (A, C, G, T, N), total coverage depth, the reference base, and per-read mapping quality (MAPQ) scores. Orphan reads were excluded, overlapping read segments were ignored to prevent double-counting, and bases were filtered using a minimum base quality of 30 and a bowtie2^54^ MAPQ of 20. Reads with a MAPQ score of 20 corresponded to 99% estimated probability of an accurate assignment per read when mapped to the species genome database. Variable sites were defined using a conservative approach that required the detection of within-mouse allelic variants (A, T, C or G) in at least 50% of experimental replicates in either HF or LF groups (i.e., at least 4 cages). This choice buffered against the possibility of sequencing or PCR errors inflating the number of variable sites tested.

For each species, nucleotide frequencies were calculated by dividing individual nucleotide counts (A, C, G, T) at each genomic position by the total coverage at that position. Per-position allele frequency changes (Δf) were calculated between the two timepoints for each subject by computing the difference in nucleotide frequency between the PRE and END timepoint for each nucleotide at each position. Within each replicate×group stratum, per-subject Δf values were averaged to form replicate-level mean changes (Δ̄f) at each site, which served as the primary input for downstream statistical testing.

Tests for allele-frequency parallelism (within diet groups) or divergence (between diet groups) were conducted on variable sites, which were defined using conservative filters designed to reduce the effects of mismapping and sequencing errors. For parallelism and divergence tests, we included positions that displayed variation in at least 50% of replicates. These position-filtering steps ensured that all sites tested exhibited adequate within-individual nucleotide variation to yield a significant difference between timepoints (for parallelism) or groups (for divergence) (based on t-tests and a *p*-value threshold of 0.05) under at least one configuration of mice-to-diet-group assignments. Consequently, the resulting percentages of variable sites per species showing significant differences between timepoints (for parallelism) or groups (for divergence) (based on t-tests and a *p*-value threshold of 0.05) were not artificially deflated by the inclusion of sites that showed insufficient variation within individual mice. These choices best allowed us to assess whether species contained a greater number of sites showing evidence of parallelism or divergence than expected under the null hypothesis assuming independence among sites (i.e., 5% of sites based on a *p*-value threshold of 0.05).

### Testing for parallel and divergent allele frequency trajectories

The following statistical testing framework was implemented in our custom AlleleFlux pipeline (https://github.com/MoellerLabPU/AlleleFlux). Multiple statistical frameworks for detecting significant allele frequency parallelism and divergence at individual genomic positions were applied independently per nucleotide (A, C, G, T) for each variable site. Tests were restricted to genomic positions (sites) detected in all eight replicates.

#### Parallelism tests

One-sample t-tests (scipy.stats.ttest_1samp, testing against a population mean of 0, two-sided) and Wilcoxon signed-rank tests (scipy.stats.wilcoxon, two-sided) were applied within each diet group independently to assess whether mean allele frequency changes at each position differed significantly from zero.

Linear mixed-effects models were fitted using statsmodels.formula.api.mixedlm with the replicate included as a random intercept. For each nucleotide at each genomic position, the allele frequency change was modeled as the response with the timepoint as a categorical fixed effect: Δ̄f ∼ time. This tested whether allele frequency changes within a single group differed significantly between timepoints while accounting for correlation among repeated measurements from the same cage/replicate. Tests were performed per nucleotide per position.

Cochran-Mantel-Haenszel (CMH) tests stratified by replicate were performed via R’s mantelhaen.test function (accessed through rpy2, two-sided, without continuity correction). Unlike the other tests the CMH test is not applied per-nucleotide; rather, it tested the joint distribution of all nucleotides simultaneously in a single contingency table per position. At each genomic position, raw nucleotide counts were aggregated per replicate within each group. Nucleotide bases with zero total counts across both groups were excluded and positions with fewer than two informative bases were skipped. Each diet group was compared between PRE and END timepoints, testing whether allele frequency distributions within that group changed over time.

#### Divergence tests

Cages from HF and LF diet groups were paired by replicate (i.e., mouse litter). Paired t-tests (scipy.stats.ttest_rel, two-sided) and Wilcoxon signed-rank tests (scipy.stats.wilcoxon, two-sided) were applied to these matched pairs.

Linear mixed-effects models were fitted using statsmodels.formula.api.mixedlm with the replicate included as a random effect. For each nucleotide at each genomic position, the mean allele frequency change was modeled as the response with the experimental group as a categorical fixed effect: Δ̄f ∼ group + 1|replicate.

CMH tests stratified by replicate were performed via R’s mantelhaen.test function (accessed through rpy2, two-sided, without continuity correction). At each genomic position, raw nucleotide counts were aggregated per replicate within each group. Nucleotide bases with zero total counts across both groups were excluded and positions with fewer than two informative bases were skipped. Using results of divergence CMH tests, we identified alleles whose frequencies differed between HF and LF cages at the END timepoint but did not differ between HF and LF cages at the PRE timepoint (referred to in Supplementary Table 2 as CMH_end_focus). We also identified alleles whose frequencies differed between HF and LF cages at the PRE timepoint but not the END timepoint (referred to in Supplementary Table 2 as CMH_pre_focus). Summary statistics based on sites whose allele frequencies differed between HF and LF cages at the END timepoint but not the PRE timepoint are shown in Extended Data Fig. 6.

#### Multiple Testing Correction

*P*-values from all statistical tests were consolidated into a unified summary table (results of significant sites at the *p* < 0.05 threshold are presented in Supplementary Data 1). For tests that produced multiple *p*-values per position (across A/C/G/T), the minimum *p*-value across the relevant sub-tests was taken as the representative *p*-value for that position and test type. For each parallelism or divergence test, Benjamini–Hochberg (BH) false discovery rate (FDR) (statsmodels.stats.multitest.multipletests) was then applied within each test type across all positions aggregated across all species. BH correction was chosen to address multiple testing across genomic positions while retaining power to detect replicated allele-frequency responses. BH controls the FDR under independence or positive regression dependence on the subset of true null hypotheses (PRDS), and thus does not require complete independence among tested sites. Because linked alleles can change frequency together through genetic hitchhiking, significant positions identify parallel or diet-associated allele-frequency responses but do not necessarily represent independent targets of selection or establish a direct fitness effect of each nucleotide.

### Diet differences in strain turnover

For each of the 62 core bacterial species, consensus average nucleotide identity (conANI) between PRE and END samples was calculated within individual mice using inStrain (v1.8.0). Within-mouse conANI values were averaged across mice within each cage, and cages with a mean conANI < 99.9% were classified as showing turnover of the dominant strain. For each species, a 2 × 2 contingency table summarized the numbers of HF and LF cages with and without strain turnover. Differences between diet groups were evaluated using permutation chi-squared tests implemented in R restricted within litter-matched pairs. P-values were corrected for multiple testing across species using the Benjamini–Hochberg (BH) method. Species with q < 0.05 were considered to show significant diet-associated differences in strain-turnover frequency, with the direction determined by the observed proportions of cages displaying turnover.

### popANI analyses and gene annotation

The reference MAGs were concatenated into a single fasta file, and the host-filtered reads were mapped back to this fasta file with bowtie2^54^ (v2.3.0). Genes were predicted with Prodigal^71^ (v2.6.3). Genes predicted by Prodigal were annotated to functional categories and protein descriptions against the Clusters of Orthologous Groups (COG20) database using reCOGnizer with default settings^72^. The bowtie2 mapping files and Prodigal gene prediction files were used as inputs for analyses in inStrain^41^ (v1.8.0). The “inStrain profile” command was used with the “--database_mode” option to generate inStrain profiles. The outputs of this command were used to run “inStrain compare” with the “--group_length 1000000 --breadth 0.05 -cov 0.025 -ani 0.95 --database_mode --store_mismatch_locations” flags. Tests for strain-replacement events based on popANI were conducted using permutation t-tests in R with package MKinfer^73^ (v1.2) with alternative = “greater” and num_permutations = 999, and method = “BH” (Benjamini-Hochberg) from base R.

### Calculating dN/dS values

The following method for calculating dN/dS values was implemented in our custom AlleleFlux pipeline (https://github.com/MoellerLabPU/AlleleFlux). We identified sites with q-value ≤ 0.05 from paired t-tests (divergence shown in Fig. 1). Positions assigned to multiple overlapping genes and positions with missing gene annotations were excluded; only sites with valid, single gene assignments were retained for downstream analysis. For each gene containing significant sites at the q-value ≤ 0.05 threshold, the ancestral sequence was reconstructed by starting with the Prodigal-predicted reference gene sequence and substituting the major allele from the PRE-timepoint at each variable position for each cage. In the case of tied allele counts, the reference base was preferred; otherwise, one was chosen randomly. For genes on the reverse strand, allele assignments were reverse complemented prior to substitution. The derived state (END timepoint) at each position was determined analogously from the END-timepoint samples from the same cage.

Sites were grouped into their respective codons based on gene coordinates and strand orientation. The Nei–Gojobori (1986) method with path averaging (NG86) was used to classify codon substitutions. For codons with k differing positions between ancestral and derived sequences, all k! possible single-nucleotide mutational pathways were enumerated. Each step along a pathway was classified as synonymous (S) or non-synonymous (N) by translating intermediate codons using NCBI genetic code table 11. Pathways passing through intermediate stop codons were excluded; terminal stop codons were permitted. The final S and N counts for each codon pair were averaged across all valid pathways. Codons where all pathways passed through intermediate stops were flagged as invalid (is_valid=False) and excluded from downstream calculations.

For each codon in the reconstructed ancestral gene sequence, the potential number of synonymous and non-synonymous sites was calculated by evaluating all 9 possible single-base changes (3 alternative bases × 3 codon positions) and classifying each as synonymous or non-synonymous. Each codon position contributed one-third of its changes to the total, yielding fractional site counts per codon. These were summed across all codons in the gene to obtain entire-gene potential S and N site counts. For each gene, observed fractional S and N counts were summed across codons, and the denominators were the entire-gene potential S and N sites (covering all codons in the gene, not only those with substitutions):

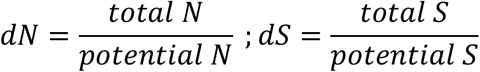

### Plotting allele frequency changes

For each statistically significant site after BH correction for MAG SLG443_DASTool_bins_SLG443_bin.96 (*q*-value ≤ 0.05, paired t-test), we identified the “anchor allele” for that position—the allele whose frequency increased over the course of the experiment in the HF group. For each site, nucleotide frequency profiles were extracted from samples belonging to the HF group at both PRE and END timepoints. Within mouse allele frequency changes were calculated for each nucleotide and then averaged across all subjects. The nucleotide with the greatest mean positive frequency change was designated as the anchor allele for that site. In the event of ties, all tied nucleotides were retained and each was tracked independently. The frequency of the anchor allele was calculated across all samples in the dataset. Samples where the coverage of the sites was below 3 reads were excluded.

Sites were ranked by minimum *p*-value, and the top 2,000 sites were selected for combined line plots. Time was represented on the x-axis as sampling day, with observations aggregated into 50-day bins (i.e., days -25–25, 25–75, 75–125, etc.); all observations with day < 0 were pooled into a single bin. The bin midpoint was used as the x-coordinate for line plots. Data points derived from fewer than 2 valid (non-NaN) samples per bin were excluded to ensure reliability of plotted trajectories. An initial frequency filter was applied to retain only sites where the mean anchor allele frequency at the first timepoint was ≤ 0.5 in both experimental groups, focusing the visualization on alleles arising or increasing during the experiment rather than those already at high frequency at baseline. The mean allele frequency per group was computed at each time bin for every site, and individual site trajectories were overlaid in a single “spaghetti plot” with lines colored by experimental group.

### Gene function enrichment analyses

To test for annotations enriched among the set of genes containing variable sites showing evidence of divergent allele-frequency changes between HF and LF groups and parallelism in the HF group, we used the annotations of all genes in the reference MAG database derived from the Clusters of Orthologous Groups COG20 database. We restricted analyses of annotation enrichment to non-mobilome genes present in bacterial species that did not show evidence of strain replacement between END and PRE timepoints in a majority of replicates based on a conANI threshold of 99.9%. For these analyses, we used hypergeometric tests in base R to test whether annotations were enriched among the set of genes containing BH-corrected significant variable sites relative to the full set of genes tested in these species. Correction for multiple testing across annotations was conducted using the BH method.

Each genomic position was mapped to its corresponding protein-coding gene using Prodigal-predicted gene coordinates. Gene boundaries were parsed from Prodigal FASTA headers (1-based, converted to 0-based coordinates) and stored in per-contig interval trees (intervaltree library https://github.com/chaimleib/intervaltree) to enable efficient positional lookups. Positions falling within annotated gene boundaries were assigned the corresponding gene identifier; intergenic positions were annotated as None/NaN.

### Analyses of nucleotide diversity within species

Nucleotide diversity (π) was quantified for the 62 core bacterial species using inStrain (v1.8.0), restricting analyses to metagenomes generated with TruSeq-equivalent library preparation from PRE, POST, and END timepoints. To standardize sequencing depth among positions, we used inStrain’s site-level rarefaction procedure with a target depth of 50 reads. Positions with fewer than 50 reads passing the inStrain filters were excluded. At each eligible position, reads were randomly subsampled to 50 before calculating nucleotide diversity. Sample-level estimates were obtained from the genome-level rarefied nucleotide-diversity output, representing the mean diversity across eligible positions in each species’ reference genome. Sample-level values were averaged within each litter, diet group, and timepoint to obtain cage-level estimates. Differences between HF and LF cages were tested separately for each species at each timepoint using two-sided Welch’s t-tests. P-values were corrected across species separately within each timepoint using the BH method.

### Bacterial cultivation and isolation

Bacterial cultivation and isolation were performed following established methods. For each dry-frozen fecal sample, 0.1g of material was diluted in 900 μl of Yeast Casitone Fatty Acids (YCFA) liquid media. 100 μl of 10^−4^ diluted samples were pipetted under anaerobic conditions onto a set of agar plates pre-reduced for ∼12 hours. All plates were incubated at 37°C for 5 days in an anaerobic (5% Hydrogen, 5% Carbon dioxide and 90% Nitrogen) chamber (Coy Lab Products Inc, MI). To isolate individual bacterial lineages, single colonies on YCFA plates were picked using toothpicks, inoculated into YCFA broth and incubated for 4–5 days in an anaerobic chamber, shaking at 37°C. Isolated bacterial cultures were then pelleted down at 1700g for 25 minutes and then stored at -80°C until used for DNA extraction.

### Isolate whole-genome sequencing and assembly

DNA extraction and library preparation for isolate whole-genome sequencing were performed following previously described methods^43^. Genomic DNA was extracted from isolates in 96-well plates using the MagMAX™ CORE Nucleic Acid Purification Kit, following the manufacturer’s instructions. For mechanical disruption, 200 µL of sample was added to 450 µL of lysis solution in cluster tubes (Axygen, cat. #MTS-11–8-C-R) containing 0.5 mm zirconia/silica beads (BioSpec, cat. #11079105z) and vortexed on a Mini-Beadbeater-96 (BioSpec, cat. #1001). Samples were disrupted for two rounds of 2.5 minutes, with a 5-minute rest between rounds. Following mechanical disruption, lysates were centrifuged for 5 minutes at 1,048 × g.

Whole-genome libraries were prepared from isolate DNA extractions using a modified Hackflex method^23,51,52^ with laboratory-made and adapted reagents from the Nextera DNA Flex Library Prep Kit (Illumina, USA). Briefly, BLT beads were diluted 1:40 in nuclease-free water (Invitrogen) and 10 µL of this dilution was mixed with 10 µL of extracted DNA and 25 µL of 2× tagmentation buffer (20 mM Tris, pH 7.6 [Invitrogen]; 20 mM MgCl₂ [Sigma]; and 50% [v/v] dimethylformamide [DMF; Sigma]). Samples were incubated at 55 °C for 15 minutes and then held at 10 °C. To stop tagmentation, 10 µL of 0.2% sodium dodecyl sulfate (SDS; Sigma) was added to each sample, followed by incubation at 37 °C for 15 minutes.

Beads were washed three times with 100 µL of washing solution consisting of a 0.22 µm MF-Millipore™ membrane-filtered solution of 10% polyethylene glycol (PEG) 8000 (Sigma) and 0.25 M NaCl (Sigma) in Tris-EDTA buffer (TE; Invitrogen). Libraries were amplified using the PrimeSTAR GXL DNA Polymerase Kit (Takara) according to the manufacturer’s protocol. Each PCR reaction contained 5 µL of 5× GXL buffer, 2 µL of 25 mM dNTPs, 1 µL of PrimeSTAR GXL polymerase, and 2 µL of nuclease-free water. The PCR mix was added directly to the washed beads, along with 5 µL of custom-synthesized i5 and i7 oligonucleotides, to a final concentration of 0.555 µM in each reaction.

PCR amplification was performed under the following conditions: 3 minutes at 68 °C; 3 minutes at 98 °C; 12 cycles of 45 seconds at 98 °C, 30 seconds at 62 °C, and 2 minutes at 68 °C; followed by 1 minute at 68 °C and a hold at 10 °C. Libraries were size selected with SPRI beads and incubated at room temperature with paramagnetic beads for 5 minutes, washed twice with 125 µL of fresh 80% ethanol, and resuspended in 35 µL of ultrapure water (Invitrogen) before transfer to a new tube.

Library concentrations and fragment size distributions were assessed using the Qubit High Sensitivity dsDNA Kit (Thermo Fisher Scientific, USA) and the High Sensitivity Bioanalyzer chip (Agilent, USA), respectively. Final isolate genome libraries were pooled in equimolar amounts and sequenced on a single P2 Illumina NextSeq 2000 flow cell at the Cornell Institute of Biotechnology. Reads were quality filtered and assembled into genomes using Bactopia v1.4 with default settings and quality controlled and assigned to taxonomic ranks using the same procedure employed for MAGs.

### Identification of invertons

We used PhaseFinder (https://github.com/XiaofangJ/PhaseFinder)^45^ with default settings to detect invertons and quantify the relative abundances of orientations in each *Bacteroides muris* and *Phocaeicola sartorii* isolate genome. The species-reference genomes of *B. muris* and *P. sartorii* assembled from wild-derived mouse metagenomes (Supplementary Table 2) were used after dereplication. Specifically, SLG221_DASTool_bins_41.fa for *P. sartorii* and SLG1207_DASTool_bins_18.fa for *B. muris* were used as references for further mapping analysis. These analyses yielded a list of confirmed invertons that displayed phase variation in reads from isolate whole-genome sequencing.

### Testing for parallel phase variation

Using the set of invertons that displayed phase variation in reads from isolate whole-genome sequencing, we tested whether each inverton reproducibly changed orientation between PRE and END in one diet group (HF or LF) but not the other diet group. For these analyses, we used PhaseFinder to calculate the proportion of reads for every inverton mapping to the forward or reverse orientation. The species-reference genomes of *B. muris* and *P. sartorii* assembled from wild-derived mouse metagenomes (Supplementary Table 2) were used as references for mapping. We then used linear mixed effects models in the R package lme4 v1.1-38 to test for a significant difference in orientation between PRE and END timepoints or PRE and POST timepoints within the HF group and within the LF group after controlling for effects of replicate. In addition, we used linear mixed effects models to test for significant differences in orientation of invertons between HF and LF mice within each timepoint after controlling for effects of replicate. Significance of differences was corrected for multiple testing across invertons using the BH method. All invertons showing BH-corrected significant differences between PRE and END timepoints in exactly one diet group were plotted in Fig. 4. No invertons showed BH-corrected significant differences between PRE and POST timepoints.

### Identification of promoters

To assess the putative functions of invertible regions, we used BPROM^50^ with default settings to predict promoter regions within invertons. For these analyses, we used BPROM to calculate linear discriminant function (LDF) scores for each promoter. Within each inverton, we classified each of the two promoters detected (each inverton only contained exactly two predicted promoters within 150bp of a gene in at least one orientation of the inverton) as ‘weak’ or ‘strong’ based on their LDF scores and their distance from the transcription start site of the closest gene. These two criteria (LDF scores and distances) yielded consistent relative rankings of ‘weak’ and ‘strong’ for the pairs of promoters identified for each inverton shown in Fig. 4. In both cases, the ‘weak’ promoter LDF score was < 5, whereas the ‘strong’ promoter LDF score was > 8, and the strong promoter was closer to the transcription start site of the nearest gene.

### Data and code availability

MAGs were generated using a previously developed snakemake workflow (github.com/CUMoellerLab/sn-mg-pipeline)^23^. Analyses of parallel and divergent allele-frequency trajectories were conducted using a custom population genetic software toolkit (https://github.com/MoellerLabPU/AlleleFlux). Inverton analyses were conducted with PhaseFinder (https://github.com/XiaofangJ/PhaseFinder)^45^. All other scripts used to conduct analyses and generate figures are made available (https://github.com/MoellerLabPU/DietDrivenMicrobiota/).

All sequencing data will be made available through the NCBI Sequence Read Archive upon acceptance of the manuscript. All genome assemblies generated from sequencing data will be made available through Dryad upon acceptance of the manuscript.

## Acknowledgements

We thank Dr. Weiwei Yan and Hannah Grazul for their help with DNA extractions, library preparations, and culturing. We also thank Anurag Agrawal, Elizabeth Johnson, and Philipp Messer for their thoughtful insight throughout the project.

## Author Contributions

A.H.M. conceived of the study idea, supervised the research, performed analyses, and wrote and edited the paper. S.G.H. conceived of the study idea, generated data, performed analyses, and wrote and edited the paper. S.U. performed analyses and wrote and edited the paper. D.M.J. performed analyses and edited the paper. A.A.L. and N.E. cultured bacteria and edited the paper. D.D.S. performed analyses and edited the paper.

## Additional Information

Supplementary information is available for this paper. Correspondence and requests for materials should be addressed to A.H.M., S.G.H., or S.U.

## Extended Data Figure and Table Legends

**Extended Data Figure 1:**
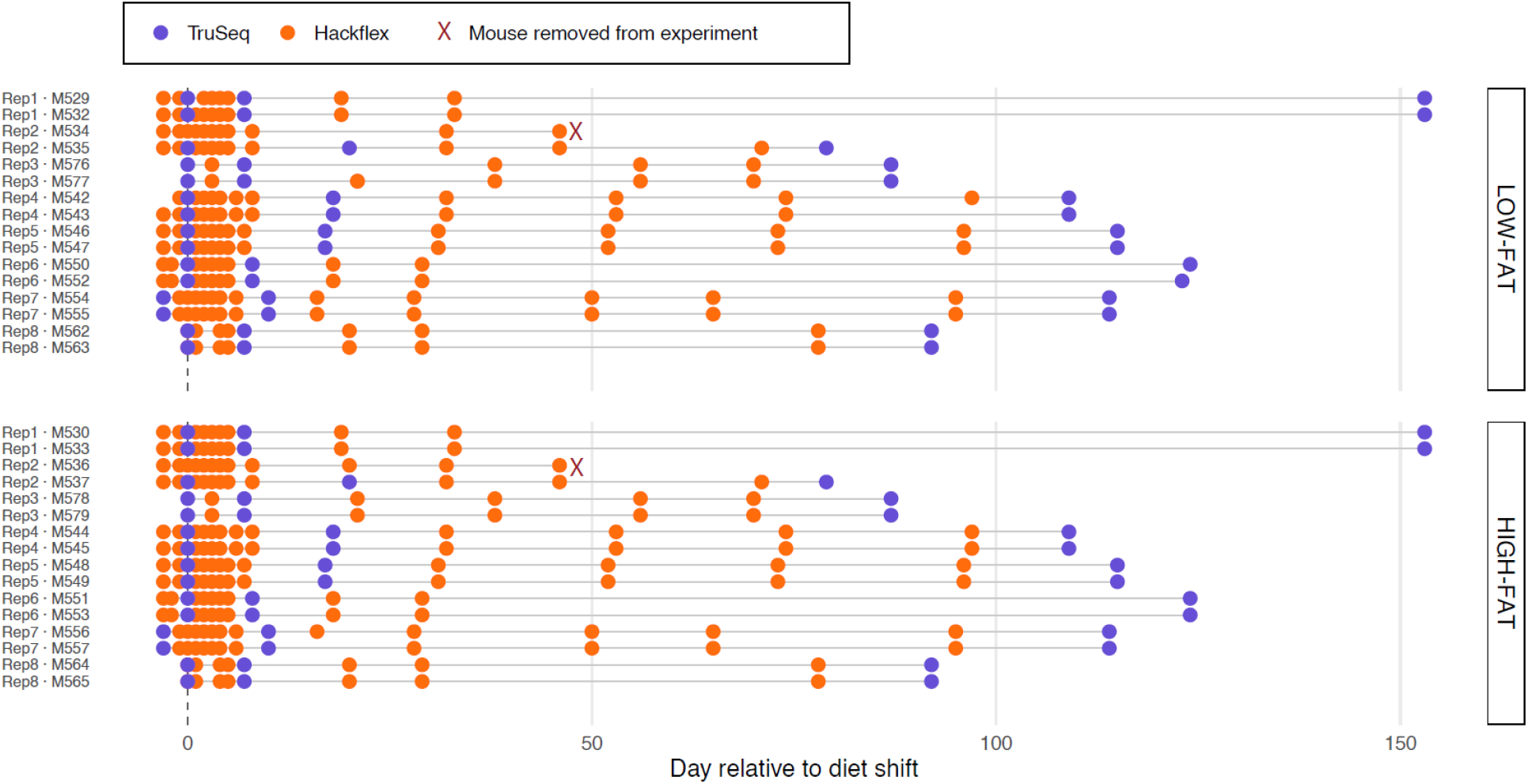
Timeline of sampling and library-preparation methods employed. Timeline shows sampling points (circles) for each mouse (lines) grouped by litter (Rep #) and treatment (LF, top; HF, bottom). Red Xs mark mice removed from the experiment due to injury. Note that within each replicate, mice were sampled on the same days, such that variation in the exact timing of sampling did not confound analyses.

**Extended Data Figure 2:**
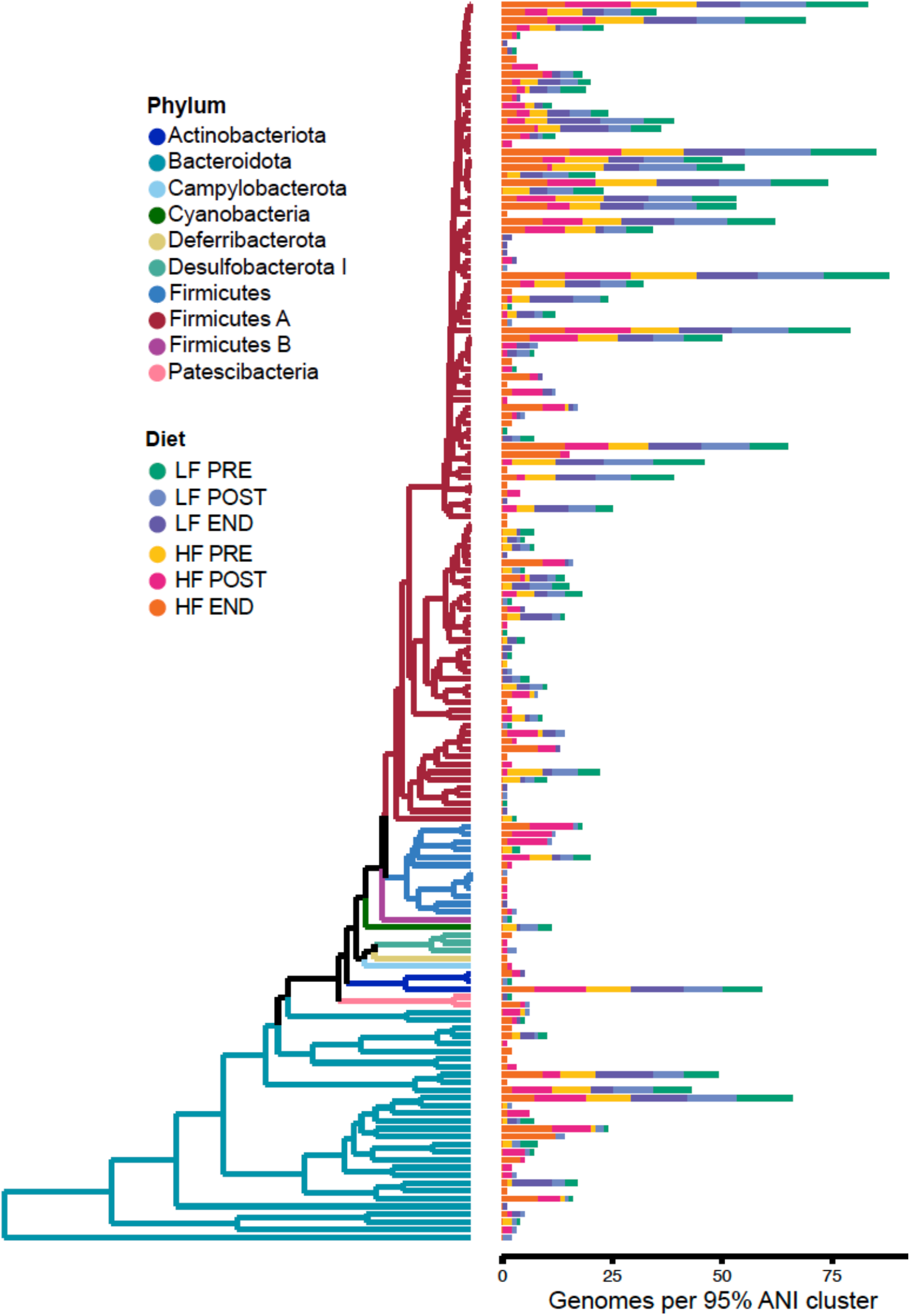
Phylogeny of MAGs. Tree shows phylogenetic relationships among 160 species representative MAGs from the reference genome database. Colors of branches denote phyla, and stacked barplots indicate the number of high-quality MAGs that form part of that species. Colors in stacked barplot indicate the diet/time group of the sample from which the MAG was assembled.

**Extended Data Figure 3:**
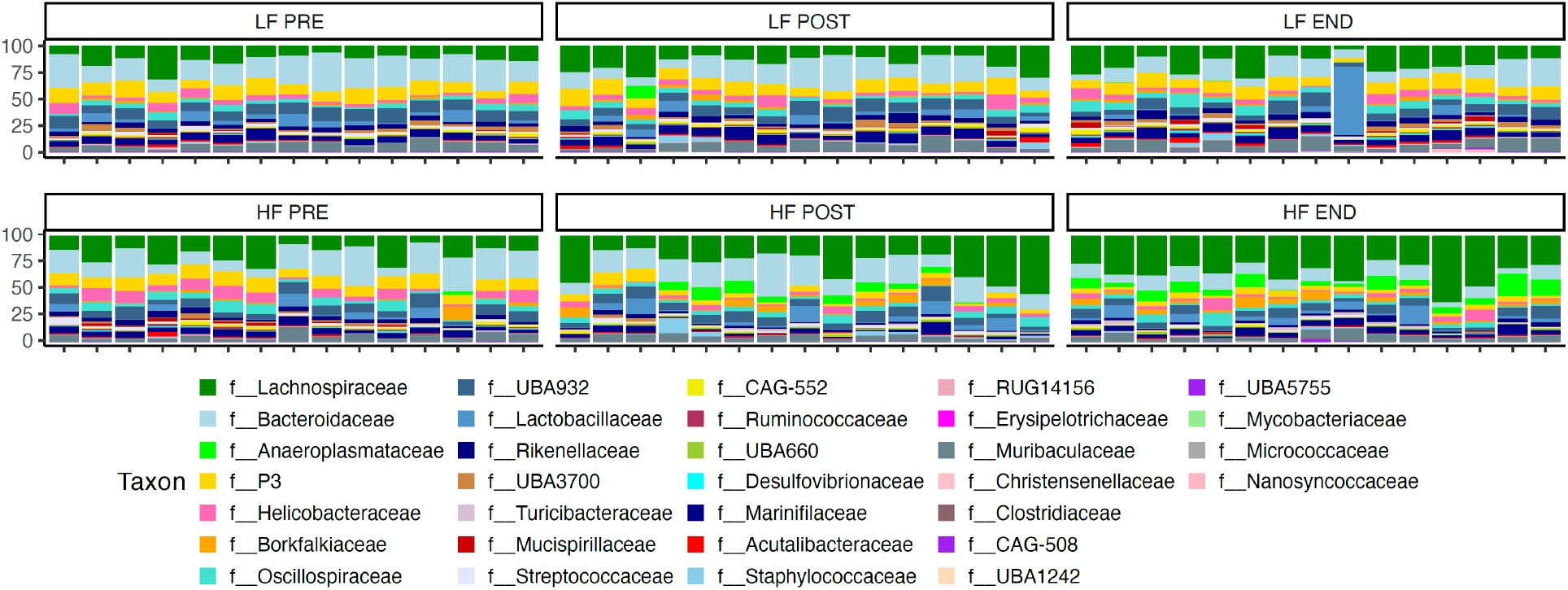
Taxonomic composition of metagenomes. Stacked bar plots show the relative abundances of bacterial families. Colors denote taxonomic assignments as indicated by the key.

**Extended Data Figure 4:**
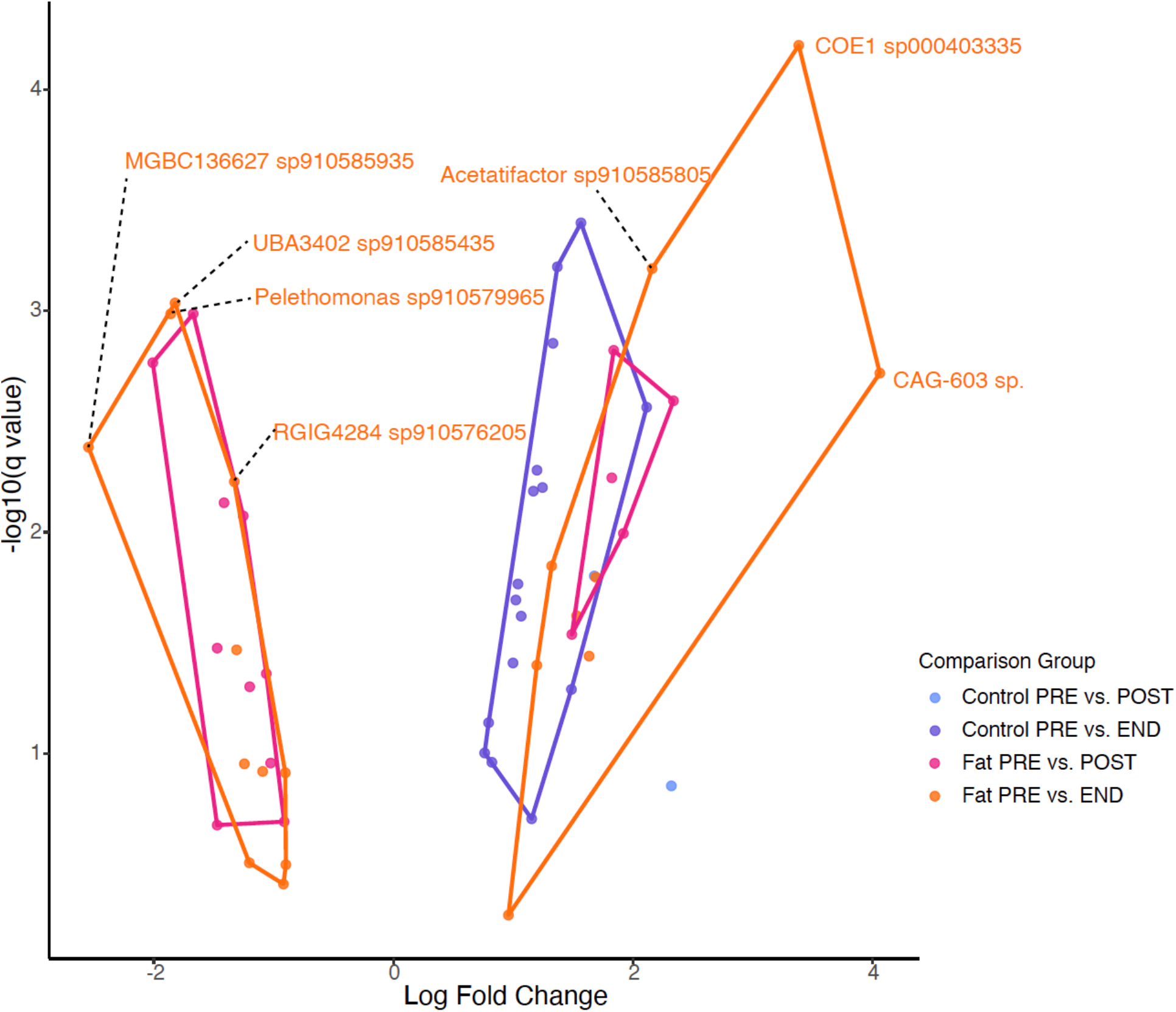
Differentially abundant taxa were most evident after the dietary transition. Volcano plots show results of ANCOM differential abundance analyses. Points denote species. Polygons enclose sets of species from the same comparisons, as indicated by the key.

**Extended Data Figure 5:**
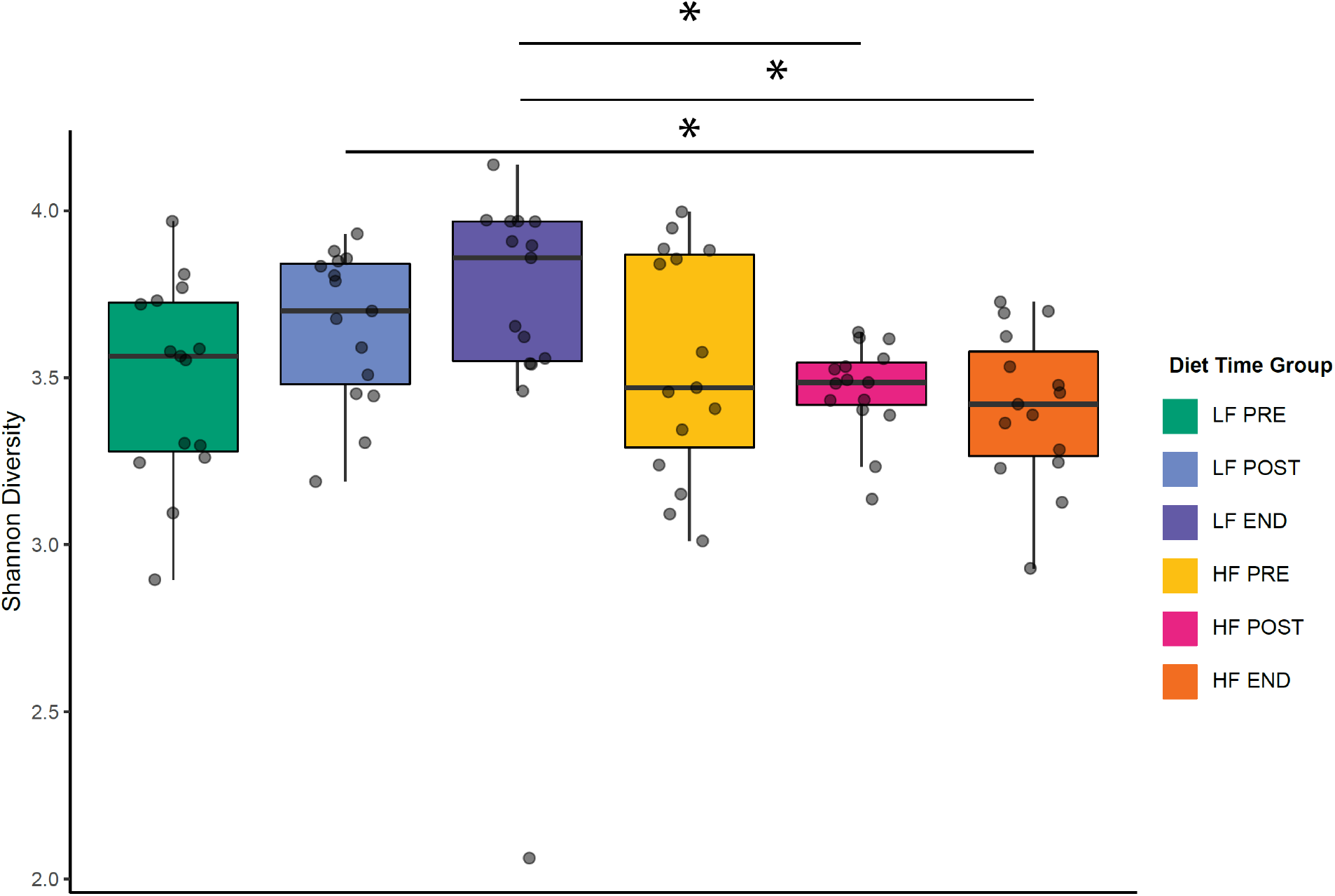
Alpha diversity differences between diet groups and timepoints. Boxplots show Shannon diversity of metagenomes. Each point is a metagenomic sample (PRE, POST, or END). Significant differences between boxplots are indicated by asterisks; Wilcoxon tests, * *p* < 0.05 ** *p* < 0.01 *** *p* < 0.001.

**Extended Data Figure 6:**
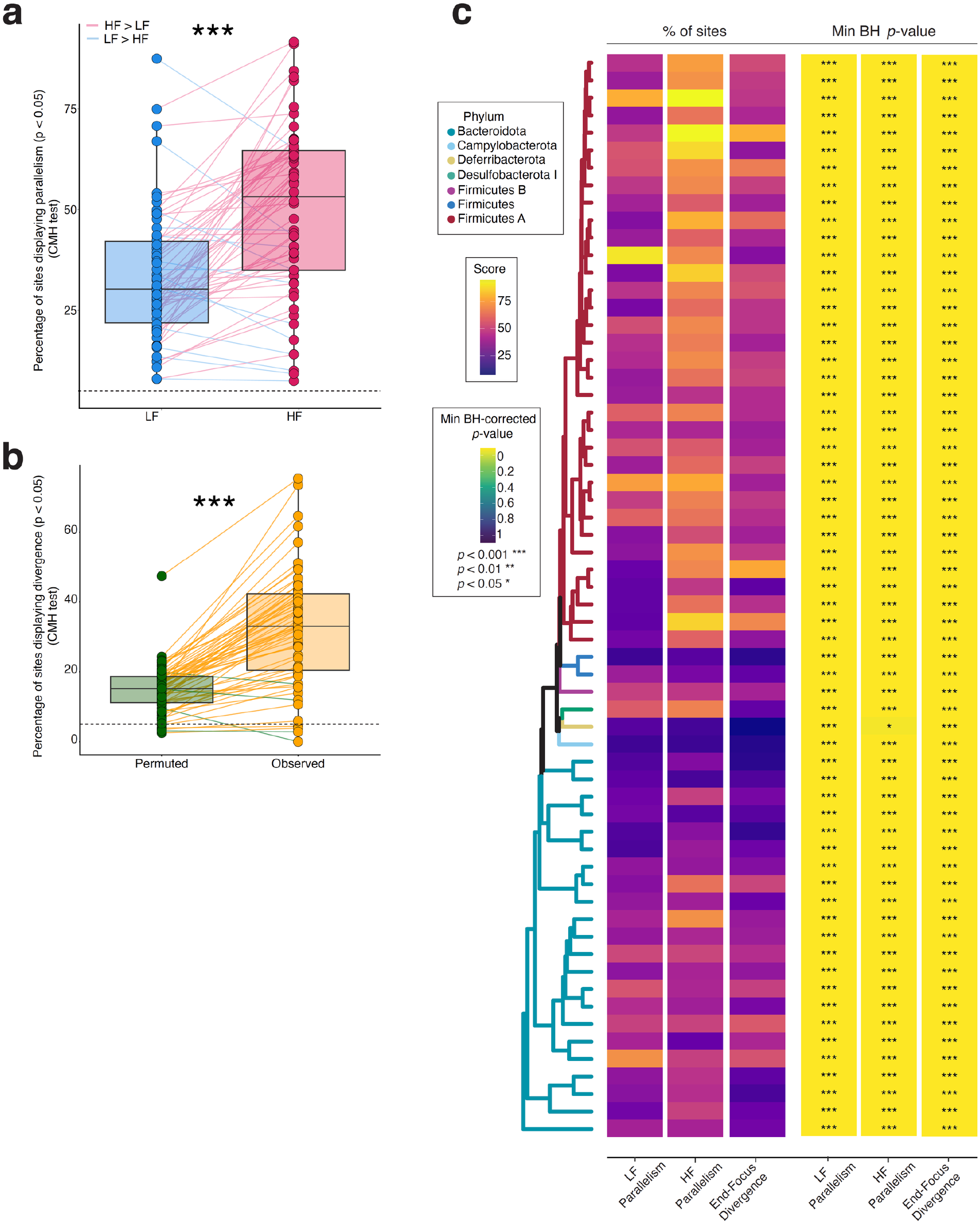
CMH tests confirm widespread adaptive responses to HF diets. a, Boxplots show the percentage of variable sites displaying parallel allele-frequency changes across replicates at the *p* < 0.05 threshold, based on CMH tests. Each point represents a bacterial species, and lines connect the same species between the LF or HF groups. Tests for parallelism compared the END to PRE samples. **b,** Boxplots show the percentage of sites displaying divergent allele-frequency changes between LF- and HF-fed mice at the *p* < 0.05 threshold, based on CMH tests. Each point represents a bacterial species, and lines connect the same species. Tests for divergence compared the changes between END and PRE samples. In **a** and **b**, asterisks indicate significance of one-sample t-test for difference in mean between boxplots; *** *p* < 0.001. **c**, Phylogeny shows relationships among 62 gut bacterial species tested for parallel and divergent allele frequency changes mediated by HF diets. Colors of branches denote phylum assignments. Leftmost three columns of heatmap show, for each species, the % of sites with *p* < 0.05 in LF parallelism, HF parallelism, or divergence tests. Rightmost three columns of heatmap show, for each species, the minimum BH-corrected *p*-value of all tested sites in that species’ genome derived from LF parallelism, HF parallelism, or divergence tests. Asterisks indicate minimum BH-corrected *p*-values; * *q* < 0.05, ** *q* < 0.01; *** *q* < 0.001. Note that the percentage of sites showing divergence in the permuted group (panel **b**) was consistently elevated above the null expectation (> 5%), indicating that CMH tests were prone to false positives.

**Extended Data Figure 7:**
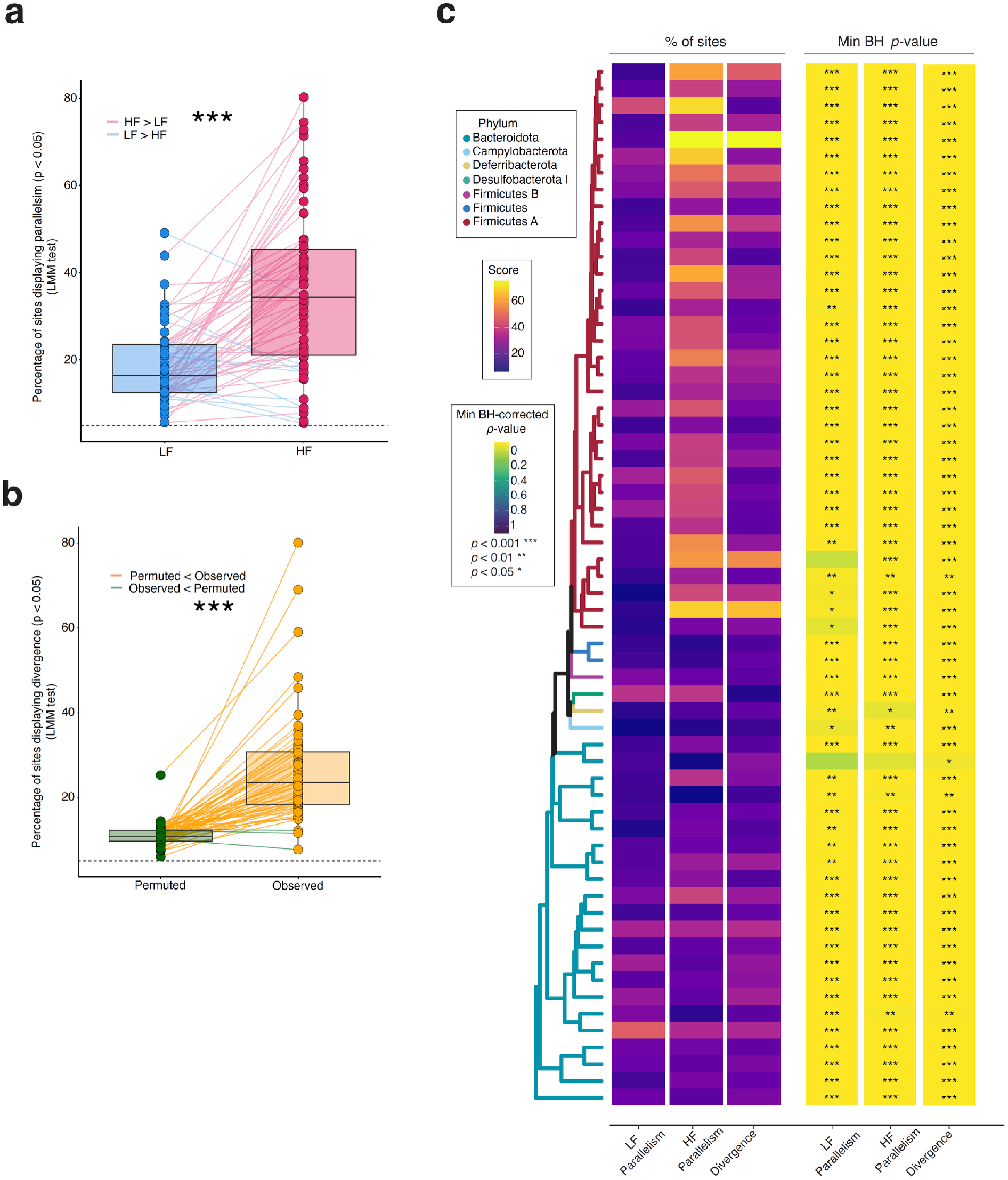
LMM tests confirm widespread adaptive responses to HF diets. a, Boxplots show the percentage of variable sites displaying parallel allele-frequency changes across replicates at the *p* < 0.05 threshold, based on LMM tests. Each point represents a bacterial species, and lines connect the same species between the LF or HF groups. Tests for parallelism compared the END to PRE samples. **b**, Boxplots show the percentage of sites displaying divergent allele-frequency changes between LF- and HF-fed mice at the *p* < 0.05 threshold, based on LMM tests. Each point represents a bacterial species, and lines connect the same species. Tests for divergence compared the changes between END and PRE samples. In **a** and **b**, asterisks indicate significance of one-sample t-test for difference in mean between boxplots; *** *p* < 0.001. **c**, Phylogeny shows relationships among 62 gut bacterial species tested for parallel and divergent allele frequency changes mediated by HF diets. Colors of branches denote phylum assignments. Leftmost three columns of heatmap show, for each species, the % of sites with *p* < 0.05 in LF parallelism, HF parallelism, or divergence tests. Rightmost three columns of heatmap show, for each species, the minimum BH-corrected *p*-value of all tested sites in that species’ genome derived from LF parallelism, HF parallelism, or divergence tests. Asterisks indicate minimum BH-corrected *p*-values; * *q* < 0.05, ** *q* < 0.01; *** *q* < 0.001. Note that the percentage of sites showing divergence in the permuted group (panel **b**) was consistently elevated above the null expectation (> 5%), indicating that LMM tests were prone to false positives.

**Extended Data Figure 8:**
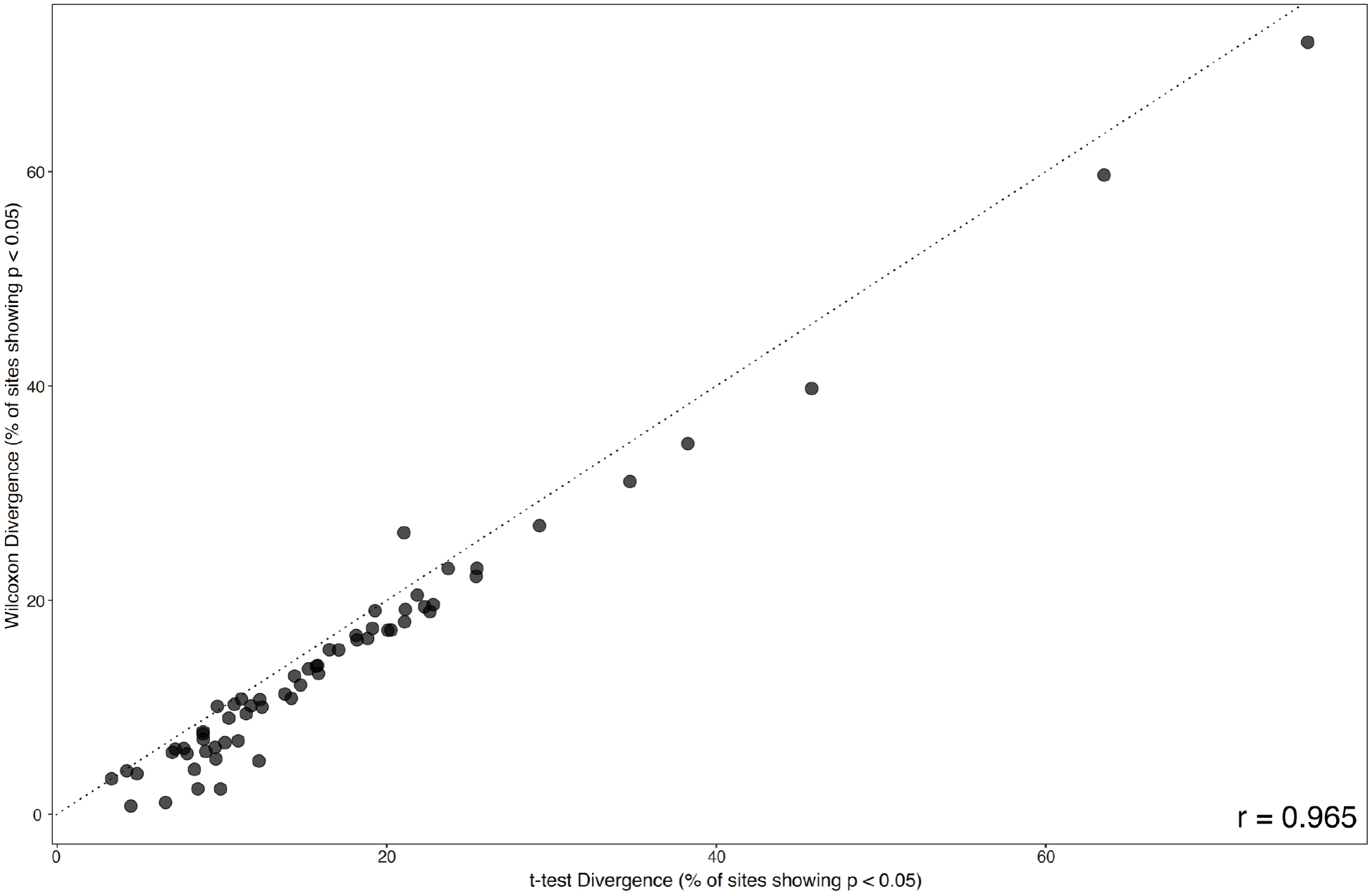
Correlation between results of parametric and non-parametric tests for allele frequency divergence. Scatterplot and regression show positive relationship between results of analyses of divergent allele frequency trajectories between HF and LF based on t-tests or non-parametric Wilcoxon tests. Each point represents a bacterial species.

**Extended Data Figure 9:**
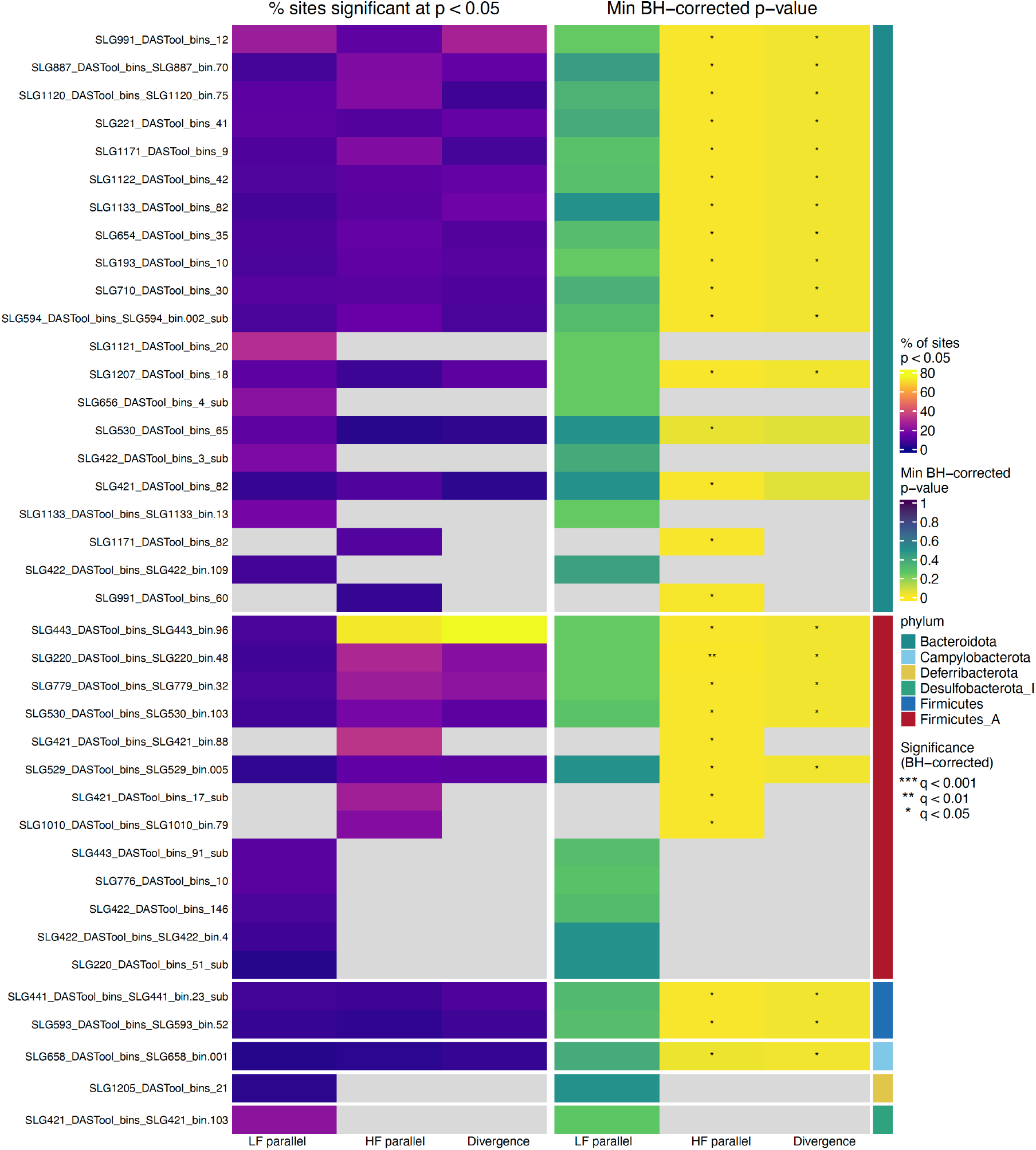
Parallel and divergent allele-frequency shifts across gut bacterial populations using a 1x coverage threshold. Heatmaps summarize allele-frequency dynamics for bacterial species (rows) during the dietary experiment. The left three columns show the percentage of variable genomic sites exhibiting significant parallel allele-frequency shifts within low-fat control (LF parallel) or high-fat (HF parallel) groups, or significant divergent allele-frequency shifts between HF and LF groups, based on a t-test *p* < 0.05 threshold. Parallelism was tested from directional mean allele-frequency shifts across replicate hosts, whereas divergence was tested from differences in allele-frequency shifts between litter-matched HF and LF mice. The right three columns show the minimum Benjamini–Hochberg (BH)-corrected *p*-value across tested sites within each MAG for the corresponding LF-parallel, HF-parallel, and divergence tests. Asterisks denote significance after BH correction: *q* < 0.05 *; *q* < 0.01 **. The colored strip at right denotes bacterial phylum. Grey cells indicate insufficient coverage for inclusion of the bacterial species.

**Extended Data Figure 10:**
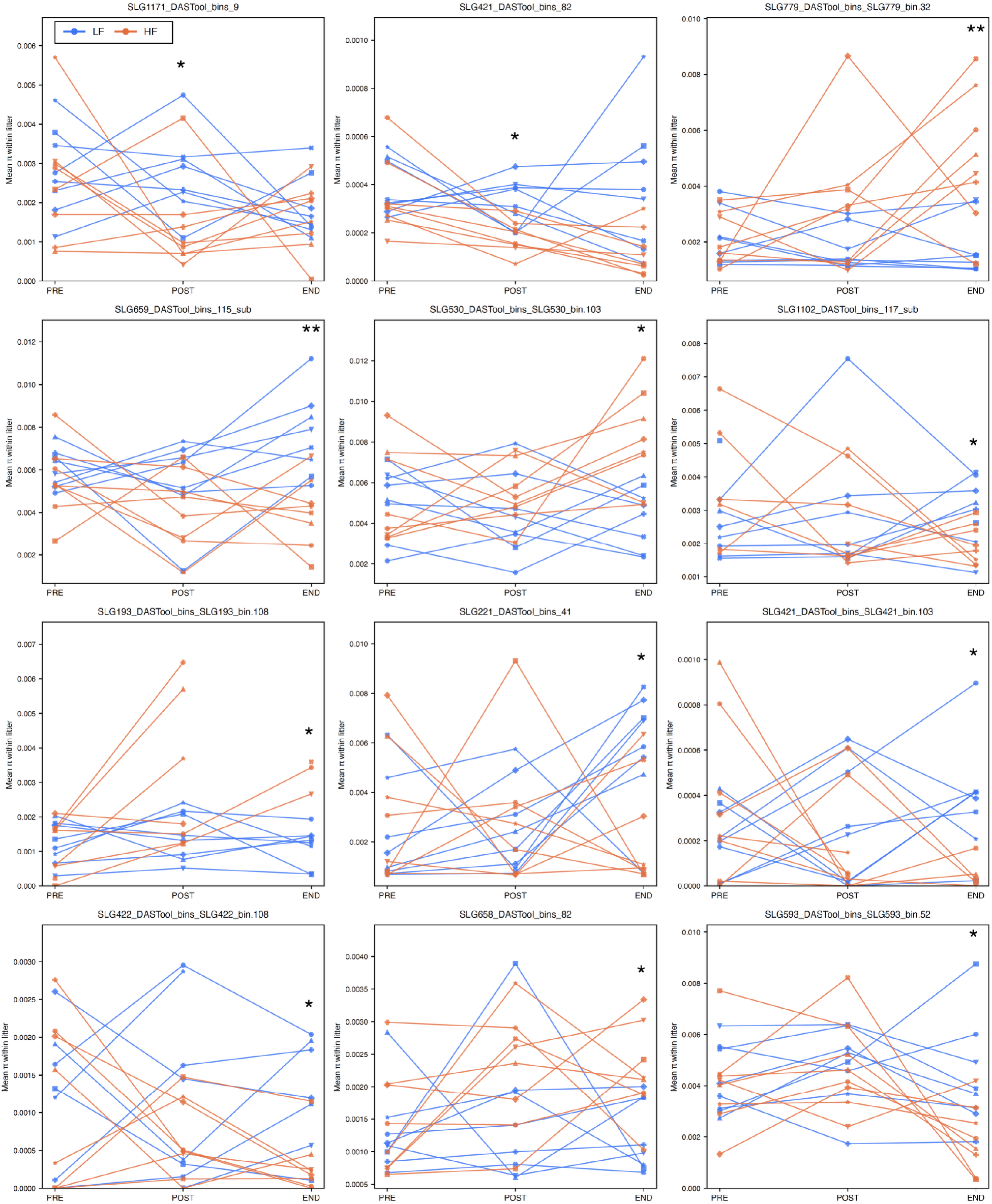
High-fat diet alters within-species nucleotide diversity in multiple gut bacterial populations. Each panel shows nucleotide diversity (π) for a bacterial MAG displaying a significant difference between high-fat (HF; orange) and low-fat control (LF; blue) groups at one or more post-treatment timepoints. π was calculated in inStrain from rarefied read sets using sites with a minimum sequencing depth of 50x. Points represent the mean π of TruSeq-sequenced samples within each litter, and lines connect litter-specific means across the PRE, POST, and END timepoints. HF and LF mice were litter-matched at the start of the experiment. Differences between HF and LF groups were tested separately for each MAG within each timepoint using two-sided Welch’s *t*-tests, with Benjamini–Hochberg correction across MAGs within each timepoint. Asterisks indicate significant differences after correction: *q* < 0.05 *; *q* < 0.01 **.

**Extended Data Table 1:** Composition of HF and LF diets. Abbreviations: LF, low-fat; HF, high-fat. Macronutrient values are given as percentage by weight or percentage of total kilocalories, as indicated. Ingredient quantities are given in g/kg diet.

| **Macronutrient composition** | **LF Diet** | **HF Diet** |
| --- | --- | --- |
| **Energy density (kcal/g)** | **3.7** | **4.2** |
| **Macronutrient % by weight** |  |  |
| **Protein** | **17.7** | **14.0** |
| **Carbohydrate** | **59.7** | **13.2** |
| **Fat** | **7.2** | **35.0** |
| **Macronutrient % kcal** |  |  |
| **Protein** | **18.9** | **13.2** |
| **Carbohydrate** | **63.8** | **12.4** |
| **Fat** | **17.3** | **74.4** |
| **Ingredient composition (g/kg)** |  |  |
| **Casein** | **200** | **158** |
| **L-Cystine** | **3** | **2.4** |
| **Corn starch** | **399.986** | **0** |
| **Maltodextrin** | **132** | **50** |
| **Sucrose** | **100** | **0** |
| **High amylose corn starch** | **0** | **110** |
| **Cocoa butter** | **0** | **258.65** |
| **Soybean oil** | **70** | **77.35** |
| **Anhydrous milkfat** | **0** | **12.42** |
| **Cellulose** | **50** | **273.13** |
| **Mineral mix, w/o Ca & P (98057)** | **0** | **13.39** |
| **Mineral mix, AIN-93G-MX (94046)** | **35** | **0** |
| **Calcium phosphate, dibasic** | **0** | **8.11** |
| **Calcium carbonate** | **0** | **6.55** |
| **Vitamin mix, Teklad (40060)** | **10** | **10** |
| **Inulin** | **0** | **10** |
| **Pectin** | **0** | **10** |
| **TBHQ (antioxidant)** | **0.014** | **0** |

