## Supplementary Information Guide for "Community-wide adaptive evolution of gut bacteria during dietary transitions"

**Supplementary Table 1.**

Fecal-sample metadata. Metadata pertaining to the fecal samples sequenced, the mice that they are derived from, and the library strategy used for sequencing. "sample" gives the unique fecal-sample identifier; "mouse" identifies the individual mouse; "replicate" gives the replicate designation; "group" indicates the dietary treatment group; "day" gives the sampling day relative to the experimental timeline; "time" gives the corresponding categorical sampling time point; "Sex" indicates mouse sex; "cage" identifies the cage in which the mouse was housed; and "library_method" indicates the sequencing-library preparation method used for the sample.

**Supplementary Table 2.**

MAG metadata. Metadata for all assembled MAGs, including quality information, genome size, classification, and results from AlleleFlux tests. "MAG_ID" gives the unique metagenome-assembled genome (MAG) identifier; "Completeness" and "Contamination" give estimated genome completeness and contamination, respectively, as percentages; "Genome Size" gives assembly size in base pairs; "High Quality" indicates whether the MAG met the criteria used for classification as high quality; "drep Representative Genome" indicates whether the MAG was selected as the representative genome of its dRep cluster; "drep Secondary Cluster" gives the dRep secondary-cluster/species-level genome-bin identifier; and "Domain", "Phylum", "Class", "Order", "Family", "Genus", and "Species" give the taxonomic classification. "percentage-of-sites-p-<-0.05_calculation" indicates whether the MAG was included in calculations of the proportion of sites with nominal $P < 0.05$. Columns beginning with "total_sites_per_group", "significant_sites_per_group", and "percentage-of-sites-p-<-0.05" give, respectively, the number of sites tested, the number with nominal $P < 0.05$, and the corresponding percentage of significant sites for the statistical analysis indicated in the suffix: "tTest", "Wilcoxon", "LMM" (linear mixed model), or "CMH" (Cochran–Mantel–Haenszel test). Additional suffixes identify the treatment or comparison to which the analysis applies, including "LF" (low-fat diet), "HF" (high-fat diet), "across_time_HF", "pre_focus", and "end_focus"; the unsuffixed paired t-test and Wilcoxon fields correspond to the paired comparison. Columns beginning with "bh_min_q" report the minimum Benjamini–Hochberg-adjusted $q$ value among sites for the indicated analysis.

**Supplementary Table 3.**

Relative abundances of bacterial species. The first column gives the MAG representing each species-level genome bin, and each subsequent column, labeled with an "SLG" identifier, represents an individual fecal sample. Values are the relative abundance of that bacterial species in the corresponding sample, expressed as a percentage; abundances across species therefore sum to approximately 100% within each sample.

**Supplementary Table 4.**

Differential abundance testing with ANCOM-BC2. Standard output from differential abundance testing from ANCOM-BC2. All testing was performed relative to the control. "Classification" gives the taxonomic classification of each bacterial species and "SGB" gives its representative species-level genome-bin/MAG identifier. For each diet group ("Control" or "Fat") and time contrast ("Post" or "End"), "Log Fold Change" gives the ANCOM-BC2 estimated natural-log fold change relative to the reference time point, "SE" gives its standard error, "W" gives the test statistic calculated from the log-fold-change estimate and its standard error, "p" gives the nominal *P* value, and "q" gives the multiple-testing-adjusted *P* value. "Different" indicates whether the taxon was called significantly differentially abundant on the basis of the specified significance threshold, and "Passed SS" indicates whether the result passed ANCOM-BC2's sensitivity analysis to the addition of different pseudocounts to zeros.

**Supplementary Table 5.**
PopANI calculations between samples. In the inStrain pairwise-comparison table, "genome" gives the reference MAG being compared, and "name1" and "name2" identify the two metagenomic samples/inStrain profiles. "coverage_overlap" is the fraction of covered positions shared between the two profiles among positions covered in either profile; "compared_bases_count" is the number of positions with sufficient coverage to be compared in both profiles; "consensus_SNPs" is the number of compared positions at which the samples have different consensus alleles; "population_SNPs" is the number at which the two samples share no called alleles; "popANI" is population average nucleotide identity, calculated from population SNPs; "conANI" is consensus average nucleotide identity, calculated from consensus SNPs; and "percent_compared" is the fraction of the reference genome that could be compared between the samples. In the strain-sorting summary at the right, "species" gives the species/SGB identifier and "species_name" its taxonomic name; "p_value_litter_0.99900" gives the Fisher's exact-test *P* value comparing the frequency of strain replacement between HF and control litters using a conANI threshold of 0.99900; "q_value_litter_0.99900" gives the corresponding multiple-testing-adjusted *q* value; "replaced_fat_litter_0.99900" and "replaced_control_litter_0.99900" give the numbers of HF and control litter units classified as undergoing strain replacement at this threshold; and "units_fat_litter_0.99900" and "units_control_litter_0.99900" give the respective numbers of evaluable litter units.

**Supplementary Table 6.** Counts and distribution of significant sites used for analyses shown in Figure 3. "species_num" gives the species/SGB cluster identifier; "species" gives the taxonomic species name; "MAG_ID" gives the representative MAG identifier; "n_sites_significant_in_both_HF-parallel_and_HF-LF-divergence" gives the number of genomic sites that were significant in both the analysis of parallel allele-frequency change in the HF treatment and the analysis of HF–LF divergence; and "n_contigs_containing_sites_significant_in_both_HF-parallel_and_HF-LF-divergence" gives the number of distinct MAG contigs containing one or more of these sites.

**Supplementary Table 7.**
Functional enrichment analyses. Results from hypergeometric tests. N = number of genes, M = number of genes in that group. "category_col" identifies the annotation field or annotation system tested for enrichment (e.g., functional category, general functional category, protein description, KEGG hit, UniRef hit, peptidase hit, or CAZy hit); "category" gives the particular annotation term being tested; "total_count" gives the number of genes/proteins in the background set assigned to that category; "significant_count" gives the number of genes/proteins in the focal significant set assigned to that category; "N" gives the total number of genes/proteins in the corresponding background universe; "M" gives the total number in the focal significant set; "hypergeo_p" gives the *P* value from the hypergeometric enrichment test; and "hypergeo_p_adj" gives the corresponding multiple-testing-adjusted *P* value.

**Supplementary Table 8.**
Results of analyses of nucleotide diversity. The summary at left uses "Time" to identify the sampling time point ("PRE", "POST", or "END") and "Result" to report the number of species/SGBs with significant differences in nucleotide diversity between HF and LF groups out of the total number tested. In the detailed results at right, "Time" gives the sampling time point, "SGB" gives the representative MAG/SGB identifier, "HF − LF π" gives the estimated difference in nucleotide diversity (π) between the high-fat and low-fat groups, with positive values indicating higher π in HF and negative values indicating higher π in LF, "p" gives the nominal *P* value for the HF–LF comparison, and "BH q" gives the corresponding Benjamini–Hochberg-adjusted *q* value.

**Supplementary Table 9.**
Metadata for isolate genome assemblies. This contains the isolate ID, assembly filename, and classification for all isolate assemblies. "isolate_ID" gives the unique identifier assigned to each cultured bacterial isolate; "assembly_filename" gives the filename of the corresponding isolate-genome assembly; and "classification" gives its complete taxonomic classification from domain through species.

**Supplementary Table 10.**
Invertons identified in isolates. The unlabeled first column is an exported row/index identifier. "ID" identifies the putative invertible DNA region using its scaffold and four boundary coordinates; "Pe_F" and "Pe_R" give the numbers of reads supporting the forward and reverse orientations, respectively, using paired-end information; "Pe_ratio" is the fraction supporting the reverse orientation, calculated as "Pe_R/(Pe_F + Pe_R)"; "Span_F" and "Span_R" give the numbers of reads directly spanning the inversion junction in the forward and reverse orientations, respectively; and "Span_ratio" is the corresponding reverse-orientation fraction, "Span_R/(Span_F + Span_R)". "isolate_ID" identifies the isolate in which the inverton was detected, and "Bacterial_species" gives the species assigned to that isolate.

**Supplementary Table 11.**
Inverton orientations in metagenomes. The unlabeled first column is an exported row/index identifier. "ID" identifies the putative invertible DNA region by scaffold and boundary coordinates; "Pe_F" and "Pe_R" give paired-end read counts supporting the forward and reverse orientations; "Pe_ratio" gives the reverse-orientation fraction calculated as "Pe_R/(Pe_F + Pe_R)"; "Span_F" and "Span_R" give the numbers of junction-spanning reads supporting the forward and reverse orientations; and "Span_ratio" gives the reverse-orientation fraction calculated as "Span_R/(Span_F + Span_R)". "MAG_ID" identifies the MAG containing the inverton, and "SLG_ID" identifies the metagenomic sample in which its orientation was quantified.

**Supplementary Data 1.**
Results of parallelism and divergence tests.

**Supplementary Results**.
Additional results from analyses not presented in the main text.
