## Supplementary Results for "Community-wide adaptive evolution of gut bacteria during dietary transitions"

Species abundance changes in response to HF diets

Differential abundance testing revealed multiple bacterial species whose relative abundances shifted in response to HF diets (Supplementary Table 4, Extended Data Figs. 2 and 3). The taxa that decreased the most in mean relative abundance in the HF group over time included Lachnospiraceae and Oscillospiraceae species, whereas other Lachnospiraceae species displayed the largest increases (COE1 sp000403335, CAG-603 sp., and *Acetatifactor* sp910585805).

Tests for divergent evolution after permuting sample-to-group assignments

Tests for divergence after randomly permuting sample-to-group assignments indicated that the approach was not biased towards false positives. In results of permuted tests for divergent allele frequency changes, the mean percentage of sites per species showing evidence of parallelism at the $p < 0.05$ threshold was not significantly different from 5% (two-tailed t-test $p > 0.9$) (Fig. 1d), as expected under the null hypothesis, indicating that the statistical approach was not biased towards false positives.

Alternative tests for parallel and divergent allele-frequency shifts

We confirmed results of t-tests using several other well-established methods for detecting significant allele-frequency shifts using data from evolve-and-resequence experiments. These analyses included non-parametric Wilcoxon tests (Supplementary Table 2), Cochran–Mantel–Haenszel (CMH) tests stratified by replicate (Extended Data Fig. 5), and tests based on linear mixed effects models (LMMs) (Extended Data Fig. 6). Results of t-tests and other tests were positively and significantly associated (Extended Data Fig. 7). In all tests for parallel evolution within HF or LF diet groups, species displayed significantly higher mean percentage of sites showing evidence of parallelism (at the $p < 0.05$ threshold) in the HF group than in the LF group. Similarly, in all tests for divergent evolution between HF and LF diet groups, species displayed significantly higher mean percentage of sites showing evidence of divergence (at the $p < 0.05$ threshold) in the real data than in the data with permuted diet-group assignments. For analyses of both parallelism and divergence, CMH and LMM approaches each detected a greater number of significant variable sites than did t-tests.

Results of popANI-based test for strain similarity across HF END cages

To test for increased strain similarity between HF END cages, we used inStrain to calculate population average nucleotide identity (popANI) between pairs of samples for every species. A popANI value close to one indicates that two populations are almost genetically identical, whereas lower values indicate greater genetic divergence. For each species, we compared popANI values between the END samples collected from HF-fed, non-littermate mice to popANI values between these samples and END samples collected from LF-fed, non-littermate mice (Supplementary Table 5). These comparisons allowed us to ask, for each species at the end of the experiment, whether the dominant strains in HF-fed mice were more similar to one another than they were to the dominant strains in LF-fed mice, after controlling for litter effects. Increased strain similarity among HF-fed mice would suggest parallel strain replacement events across HF-fed mice from different litters. Based on permutation t-tests, we found that ~48% (30/62) of species displayed significantly higher popANI between HF-fed mice than between HF- and LF-fed mice (BH-corrected $p < 0.05$) (Supplementary Table 5), indicative of parallel strain replacements mediated by HF diets. Compared to other species, these species showing popANI clustering had significantly higher mean percentage of sites showing allele-frequency divergence at the $p < 0.05$ threshold (based on t-tests) between HF and LF groups (one-sided t-test, $p = 0.046$). Moreover, across species, the test statistics of the permutation t-tests for strain replacements were positively associated with the percentage of species' variable sites in showing evidence of allele-frequency parallelism at the $p < 0.05$ threshold (based on t-tests) in the HF diet (test statistic vs. HF parallelism %; Spearman's rho = 0.356, $p = 0.0057$). The test statistics of the permutation t-tests for strain replacement were also positively associated with the percentage of sites showing evidence of allele-frequency divergence at the $p < 0.05$ threshold (based on t-tests) (test statistic vs. divergence %; Spearman's rho = 0.49, $p$ = 7.553e-5). The Lachnospiraceae species shown in Fig. 2 displayed the strongest evidence for strain replacement (BH-corrected permutation t-test $p$ = 2.204e-7).

<u>Strain replacement of Lachnospiraceae species mediated by HF diets</u>
A Lachnospiraceae species (UBA3282 sp009774655) displayed the clearest evidence of HF diet–mediated parallel strain replacement. This species also had the greatest number of significant BH-corrected divergence and HF parallelism $p$-values (Supplementary Dataset 1), corresponding ~64% (62128 of 97604) and ~63% (64011 of 102154) of variable sites for the divergence and HF parallelism tests, respectively. In contrast, none of the variable sites in this species displayed significant parallel changes in allele frequencies in the control group after BH correction.

<u>Validation of allele-frequency shifts detected in metagenomic data using isolate data</u>
We used the genomes from *P. sartorii* and *B. muris* to confirm HF-induced changes in allele frequencies detected by analyses of metagenomic data. For each variable position in the *P. sartorii* and *B. muris* genomes detected in metagenomic data and tested for divergent allele-frequency trajectories between diet groups, we determined the major allele present in isolates derived from the END and PRE samples collected from HF and LF groups. We then tested whether the nucleotide present at these sites in metagenomes and isolates from the same diet group and timepoint agreed or disagreed. These analyses confirmed that the major alleles detected by isolate whole-genome sequencing and metagenomic sequencing were identical for > 99.9% of variable positions for *P. sartorii* and > 99.9% of variable positions for *B. muris*. These results confirm adaptive responses of these species following HF dietary transitions

<u>Confirmation of inverton phase changes using allele-frequency estimates</u>
The genome of *Phocaeicola sartorii* contained two invertons whose orientation significantly changed between END and PRE timepoints in one diet group but not the other group. Both invertons were within 100 kilobases in the *P. sartorii* genome and significantly reversed orientation across replicates after the transition to the HF diet but did not change orientation in the LF-diet group (Fig. 4a). These invertons also contained positions that showed significantly divergent allele-frequency trajectories between HF- and LF-diet groups (Fig. 1e; Supplementary Dataset 1), and each of these positions was within one sequencing-read length of a palindrome flanking the inverton, as expected if the inverton orientations reversed in response to HF diets. Moreover, in every case, the specific nucleotides within these invertons displaying divergent allele-frequency trajectories between diet groups (Fig. 1e; Supplementary Dataset 1) matched the nucleotide expected if the inverton had reversed orientation.

Functional consequences of HF-induced invertons in P. sartorii

For the inverton spanning positions 209006–209503 in the *P. sartorii* reference genome (Supplementary Table 3), we observed tandem homologs of *susC* and *susD* in the 5' flanking region with a start codon indicating that the invertible region was upstream of transcription of these genes (Fig. 4). The 3' flanking region did not contain protein-coding genes. The inverton contained two promoter sequences within 150 base pairs (bp) of the transcription start site of the 5' flanking *susC*/*susD* genes in at least one orientation of the inverton (Fig. 4b). These promoters differed in their strength based on BPROM linear discriminant function scores[42], with the reverse orientation corresponding to relatively stronger promotion of expression of *susC*/*susD* and the forward orientation corresponding to relatively weaker promotion of expression of these genes. Notably, the invertase motif separating the 5' flanking region and the invertible region contained a specific sequence (AGTTC) previously linked to the expression of *susC*—a gene that codes for a cell-surface glycan foraging protein—in *Bacteroides fragilis,* another species within the Bacteroidaceae[74].

For the inverton spanning positions 108998–109206 in the *P. sartorii* reference genome (Supplementary Table 2), we observed five open reading frames annotated as genes of unknown function in tandem in the flanking 5' region with a start codon indicating that the inverton was upstream of transcription of these genes. The 3' flanking region did not contain any protein-coding genes. The inverton contained two promoter sequences within 150 base pairs (bp) of the transcription start site of the 5' flanking genes in at least one orientation of the inverton (Fig. 4c). These promoters differed in their strength based on BPROM linear discriminant function scores[50], with the reverse orientation corresponding to relatively stronger promotion of expression of the genes and the forward orientation corresponding to relative weaker promotion of expression of these genes, suggesting that the inversion induced by HF-diets tuned expression of these genes of unknown function in ways that paralleled the inverton upstream of *susC* and *susD*.
